# PD-1 blockade enhances T cell activation by reorchestrating CD28 and CTLA4 ligand interactions

**DOI:** 10.64898/2026.04.12.718055

**Authors:** Gee Sung Eun, Jonathan Boyd, Vinh Au, Andrew Garcia, Saso Cemerski, Simon J. Dovedi, Kristen N. Pollizzi, Michael L. Dustin, Rajat Varma

**Affiliations:** Oncology R&D, AstraZeneca, Gaithersburg, MD, USA; Discovery Sciences, Biopharmaceuticals R&D, AstraZeneca, Gaithersburg, MD, USA; Biologics Engineering, R&D, AstraZeneca, Gaithersburg, MD, USA; Oncology R&D, AstraZeneca, Cambridge, UK; The Kennedy Institute and The Chinese Academy of Medical Sciences Oxford Institute, University of Oxford, Oxford, UK

## Abstract

The network of interactions comprising CD28, CTLA4 and PD-1 and their ligands CD80, CD86, and PD-L1 have profound implications for T cell activation. Despite their critical role, factors that determine their receptor occupancy when all three ligands are present remains incompletely understood. Using a supported lipid bilayer to replicate the antigen-presenting cell membrane facilitated a quantitative analysis of CD80, CD86, and PD-L1 interactions. Our observations reveal that PD-1 blockade operates through a previously unrecognized competitive mechanism beyond conventional checkpoint inhibition: liberated PD-L1 molecules redistribute to form CD80-PD-L1 heterodimers that strengthen CD28-CD80 interactions and outcompete CD86 for CD28 binding. This preferential binding enhances T cell activation through superior CD80 co-stimulation. Furthermore, the redistribution by PD-1 blockade reduces CD80 homodimer availability, limiting CTLA4 binding to CD80 and enhancing CD28 signaling. These findings provide vital insights into the competitive principles that allow B7-CD28 family receptors to regulate T cell activation through network-level effects.

## Introduction

The two signal hypothesis for T cell activation states that stimulating T cells through the T cell receptor (TCR) alone leads to anergy or tolerance, but the addition of a second signal through a co-stimulatory receptor such as CD28 would lead to complete T cell activation (*1*). Since then, multiple co-stimulatory and co-inhibitory molecules and their ligands that regulate and fine tune immune responses have been discovered (*2, 3*). Among these, the B7-CD28 family plays an important role in regulating immune responses and has been the target of therapies for cancer and autoimmune diseases (*4–6*). CD28 has two ligands: CD80 and CD86 (*7, 8*). In the 1990s, another receptor of the B7 family was discovered which was named cytotoxic T lymphocyte antigen 4 (CTLA4), which shares its ligands with CD28 and was shown to be an inhibitory receptor (*9, 10*). CTLA4 binds to CD80 and CD86 at higher affinities than CD28 and binds to CD80 at higher affinity than CD86 (*11, 12*), setting up the following hierarchy of binding interactions CTLA4-CD80 > CTLA4-CD86 > CD28-CD80 > CD28-CD86 (*13*). Subsequently, another inhibitory B7-CD28 family member, programmed cell death 1 receptor (PD-1) was discovered (*14*). PD-1 has two ligands, PD-L1 and PD-L2 and it binds to PD-L2 with higher affinity than PD-L1 (*15, 16*).

Mice deficient in CD28 show impaired T cell responses (*17*), while mice deficient in CTLA4 develop a lymphoproliferative disorder that is mirrored in humans, establishing it as an inhibitory receptor (*18–20*). CD28 deficient humans, on the other hand, are generally healthy but are susceptible to skin papillomavirus infection (*21*). The expressions of CD28 and CTLA4 are also highly regulated upon T cell activation. CTLA4 is either very low or absent on naïve T cells and is upregulated upon activation, while CD28 is expressed on naïve T cells and its expression is further increased on activated T cells (*22–24*).

The expression of PD-1 is absent on naïve mouse T cells and can be present at very low levels on human naïve T cells. PD-1 expression is upregulated upon T cell activation, and its expression is a hallmark of dysfunctional T cells in cancer and chronic infections (*25, 26*). Both PD-L1 and PD-L2 are largely expressed on myeloid cells and their expression can also be found on fibroblasts, trophoblasts and tumors (*27, 28*). The inhibitory effects of PD-1 are mediated through recruitment of the phosphatase SHP-2 to the cytoplasmic tail, that impacts the phosphorylation of CD28 and other signaling molecules (*29–31*). Multiple antibodies targeting these checkpoints are approved as cancer immunotherapies (*32–34*).

CTLA4 acts as an inhibitor of CD28 largely through competition for its ligands (*9, 10*). Both CTLA4 and CD28 exist as covalent, disulfide-linked dimers on the cell surface (*35, 36*). CD80, and less so CD86, exist as a low-affinity non-covalent homodimers through interactions in the extracellular domains, thus creating a pool of dimers and monomers (*37, 38*). Each CTLA4 dimer can interact with two CD80 or CD86 monomers or homodimers (primarily CD80) and forms chain like lattices with CD80 dimers, at least in crystal lattices, while CD28 homodimers can only interact with one CD80 or CD86 at a time (*39–41*). CD80 and CD86 cannot form heterodimers (*42*). PD-1, PD-L1 and PD-L2 all form weak homodimers through transmembrane domain interactions (*42*). While the impact of the PD-L1 and PD-L2 homodimers has not been studied, eliminating PD-1 homodimerization reduces its inhibitory potential (*42*). The discovery that CD80 and PD-L1 form heterodimers in *cis,* based on extracellular domain interactions, established a potential for a crosstalk between the PD-1 pathway and the CD28/CTLA4 pathway (*43, 44*). CD80-PD-L1 *cis-*heterodimers have reduced binding to PD-1 (*44*). CD80-PD-L1 heterodimers can still interact with CTLA4 but may not form chain like interactions that can be formed with CD80 homodimers (*44–46*). These findings add another layer of complexity to signaling within the B7 family that may influence therapeutic outcomes of anti-PD-1/PD-L1 therapies. Lastly, both CD80 and CD86 have been shown to be trans-endocytosed by CTLA4 (*47*), however, CD86 interactions with CTLA4 can be released at lower pH causing recycling of CTLA4 (*48*). In the context of CD80-PD-L1 heterodimers, only CD80 is trans-endocytosed and PD-L1 remains on the cell surface making it available for interactions with PD-1 (*48*). While the trans-endocytosed of CD80 adds another layer of complexity in this network of interactions, these effects would cause immunomodulation at longer time scales (hours).

A predictive understanding of how CD28, CTLA4 and PD-1 modulate immune responses in naïve and activated T cells when all the three ligands, CD80, CD86 and PD-L1 are present remains elusive. Furthermore, receptor occupancy of CD28, CTLA4 and PD-1 by their respective ligands remain poorly understood when these ligands are present at physiological levels and can undergo homo- and hetero-dimerization. Existing studies that have analyzed receptor-ligand interactions in isolation fail to capture the full extent of competing interactions that can occur in this pathway. We sought to address these questions by quantifying the accumulation of CD80, CD86 and PD-L1 in the immune synapse formed between a T cell and a glass supported lipid bilayer that contains these ligands, adhesion molecules and anti-TCR antibodies and correlating ligand accumulation with T cell activation. We performed PD-1 and CTLA4 blockade and competition experiments to understand how ligands switch between receptors. Our results provide mechanistic insights into how PD-1 blockade enhances T cell activation and the limits of CD80/PD-L1 *cis*-heterodimers to block PD-1 binding and CTLA4 binding.

## Results

### A microscopy-based system to study protein-protein interactions in the CD28/CTLA4/PD-1 pathway at the immunological synapse formed by human T cells

The glass supported lipid bilayer system has been extensively used to study two-dimensional protein-protein interactions at the immune synapse formed by T cells (*49, 50*). The flat geometry allows for high resolution quantitative microscopy. In addition, it offers a reductionist approach through the ability to modulate the type and quantities of molecules incorporated in the bilayer. Proteins in the lipid bilayer bound to the receptors on the cell surface appear clustered in the contact interface and their pattern reflects the spatial pattern of the receptor-ligand interactions (*51*). To study protein-protein interactions in the CD28/CTLA4/PD-1 pathway using human T cells, we reconstituted supported lipid bilayers (SLBs) with anti-TCR variable heavy domain of the heavy chain antibodies (VHH), and extracellular domain (ECD) of ICAM-1 (fig. S1A). We assessed the quality of SLBs by measuring the lateral mobility of ICAM-1 attached on the SLBs using fluorescence recovery after photobleaching (FRAP) (fig. S1B) and determined the density of incorporated molecules, such as PD-L1, using single-molecule approaches. (fig. S1C). The his-tagged anti-TCR VHH incorporated in the SLB promoted immunological synapse formation by primary human T cells from healthy donors and was associated with T cell adhesion and clustering of AlexaFluor^®^(AF)-555 labeled ICAM-1^ECD^-His (fig. S1D).

### CD28 interactions with CD86 persists in the presence of CD80 or CD80-PD-L1 at the immune synapse

Employing this SLB system, we first examined CD28-ligand interactions in two distinct SLBs composed with either ECD of fluorescently labeled CD80-His or CD86-His, in addition to anti-TCR VHH-His and ICAM-1^ECD^-His. Consistent with earlier studies and higher affinity of CD28 for CD80 than CD86 (*11*), CD28-CD80 formed stronger interactions than CD28-CD86. This was evident by stronger ligand cluster signals, measured as CD80- or CD86-AF555 intensity fold change at the immune synapse (Fig. 1A). These comparisons were performed separately using freshly isolated primary CD3^+^ T cells from healthy donors. While these were not sorted for naïve T cells, we expect CTLA4 levels to be very low in these cells. To understand how CD80 and CD86 compete for CD28 binding, we employed the Jurkat T cell line which expresses higher levels of membrane CD28 compared to primary human CD3^+^ T cells and has no expression of CTLA4. Consistent with results obtained with primary T cells, we found that CD28 bound to CD80 more strongly than CD86. When Jurkat T cells were incubated on bilayers that contained both CD80 and CD86 at a ratio of 1:1, the accumulation of CD86 decreased significantly suggesting that CD80 outcompeted CD86 for binding CD28, though not completely (Fig. 1B, ‘- PD-L1’). We next tested different ratios of CD80:CD86 to find conditions under which we may observe equivalent binding of these two ligands to CD28. While CD80 dominated at 1:1 ratio, a ∼9-fold excess of CD86 was needed to achieve comparable clustering to CD80 (Fig. 1C). Since human APCs express higher levels of CD86 than CD80 (*52*), these ratios may reflect physiologically relevant conditions. Our quantitative imaging data demonstrates that despite weaker CD28-CD86 affinity (*11*), an excess of CD86 can balance receptor occupancy of CD28 by CD80 and CD86 at the immune synapse, setting the stage for investigating effects of checkpoint blockade.

**Fig. 1.**
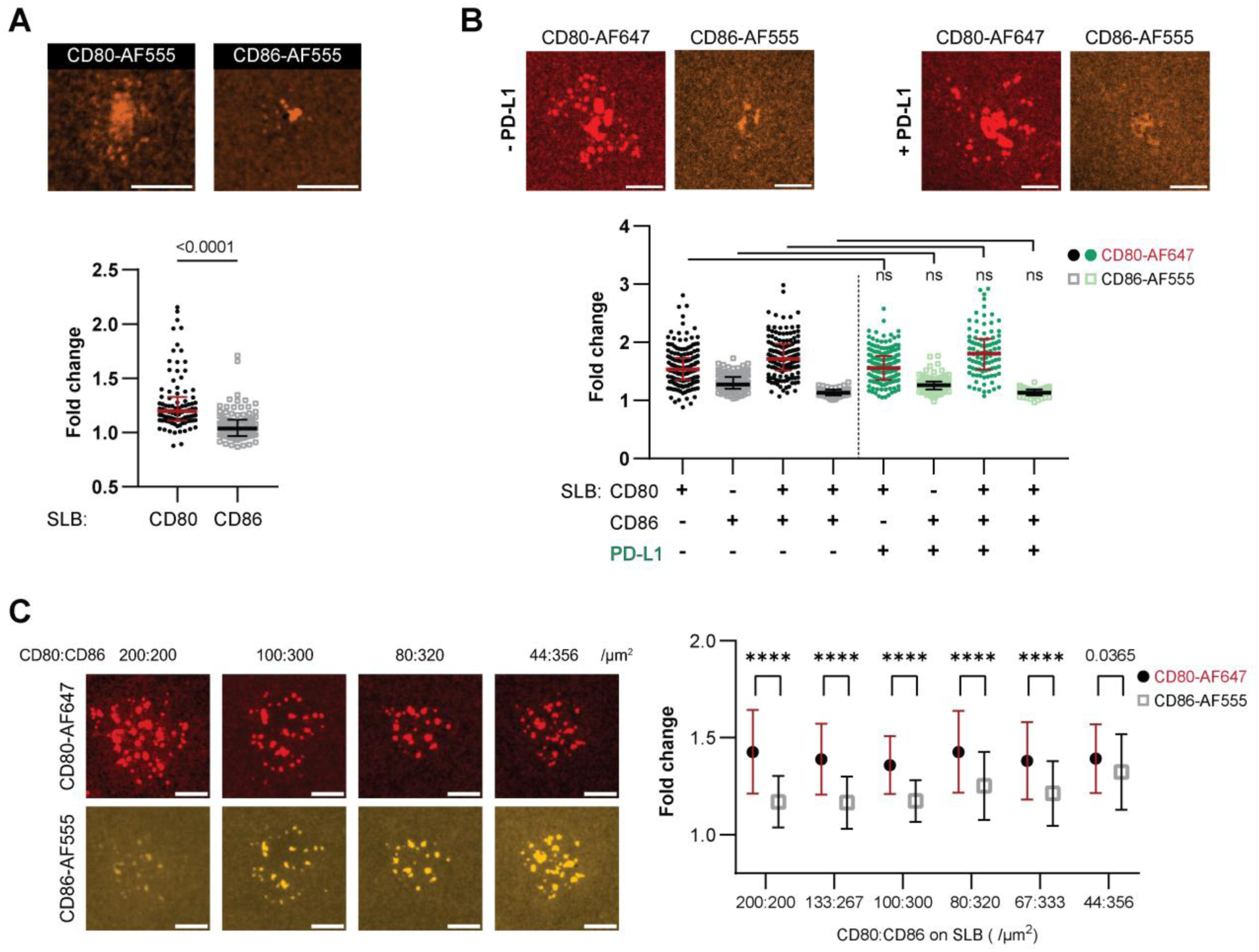
CD28 interactions with CD86 persists in the presence of CD80 or CD80-PD-L1 at the immune synapse. **(A)** Immune synapses formed by freshly isolated human primary CD3^+^ T cells on supported lipid bilayers (SLBs) containing anti-TCR VHH, ICAM-1, with either CD80-AF555 or CD86-AF555 were imaged using spinning disc confocal microscopy. Upper panel: Representative images of immune synapses on bilayers containing CD80 (left panel) or CD86 (right panel) are shown. Scale bars: 5 µm. Lower panel: Quantification of CD80 and CD86 accumulation per synapse as a fold change over area of bilayers with no cells from either CD80-AF555 SLB or CD86-AF555 SLB. Data presented as a dot column plot with median ± interquartile range (N_Cells_ >90, from two independent experiments). P value determined by unpaired t-test. **(B)** Immune synapses formed by Jurkat T cells on SLBs containing anti-TCR VHH, ICAM-1, CD80-AF647, CD86-AF555, with or without PD-L1 were imaged using spinning disc confocal microscopy. Upper panel: Representative images showing the accumulation of CD80 or CD86 in the absence (left two panels) or presence (right two panels) of PD-L1. Scale bars: 5 µm. Lower panel: Quantification of CD80 (circles, odd columns) and CD86 (squares, even columns) accumulation per synapse as a fold change over area of bilayer with no cell and the impact of PD-L1 is shown. Data presented as dot column plots with median ± interquartile range (N_Cells_ > 150, from three independent experiments). Unpaired t-tests were performed to investigate the impact of PD-L1 on CD80 or CD86 accumulation. **(C)** CD80 and CD86 competition for CD28 at different ratios. Jurkat T cells forming immune synapses on SLBs containing anti-TCR VHH, ICAM-1, and different ratios of CD80-AF647 and CD86-AF555 were imaged using spinning disc confocal microscopy. The total of CD80 and CD86 densities were maintained constant. Left panel: Representative images showing fold accumulation of CD80 (top panels) and CD86 (lower panels) from four different ratios of CD80:CD86 are shown. Scale bars: 5 µm. Right panel: Quantification of CD80 versus CD86 fold accumulation at different ratios. Data is presented as mean ± SD (N_Cells_ >250, from three independent experiments). P values determined by unpaired t-tests; ****, p<0.0001.

We next investigated whether CD80-PD-L1 *cis-*interactions (*43, 44*) affect the affinity of CD28 for CD80 and therefore the competition between CD80 and CD86 for CD28, in a system where no other receptors of interest (CTLA4 and PD-1) exist. We first confirmed that we can observe CD80-PD-L1 *cis*-interactions in the immune synapses using the SLB system (fig. S2). Using freshly isolated T cells and Jurkat T cells that express minimal PD-1, we found that fluorescently labeled PD-L1 clustered at the immune synapse only when CD80 was present (fig. S2A). This PD-L1 clustering decreased in the presence of anti-PD-L1 blockade (fig. S2B) as the site on PD-L1 that interacts with CD80 is overlapping with the one that interacts with PD-1 (*43, 53*).

To test whether the *cis-*interaction of CD80-PD-L1 influences the competition of CD80 and CD86 interaction to CD28, we added PD-L1^ECD^-His to SLB and compared fluorescently labeled CD80 cluster intensities and CD86 cluster intensities. The addition of PD-L1 did not change CD28-CD80 and CD28-CD86 interactions (Fig 1B, ‘CD80/CD86/PD-L1(+/+/+)’). These results show PD-L1-CD80 *cis*-interaction did not alter the affinity of CD28-CD80 interactions.

### CTLA4 leads to increased interactions between CD28 and CD86 due to its higher affinity to CD80

We next investigated how CTLA4, the checkpoint receptor that shares its ligands with CD28 competes with CD28 for CD80 and CD86. As introduced, previous studies have established a clear affinity hierarchy among [CD28, CTLA4] × [CD80, CD86] interactions, with CTLA4-CD80 representing the strongest binding pair (*11*), and its preference for CD80 homodimers adds an additional avidity component (*38*). To assess how CTLA4 competes with CD28 for binding to CD80 or CD86, we performed experiments with Jurkat T cells that express CD28 and no CTLA4 (fig. S3A) and tested the ability of a soluble recombinant CTLA4-Ig fusion protein to compete with CD28 for CD80 and CD86 in the lipid bilayer (Fig. 2). To validate this assay, we tested Jurkat T cells on SLBs containing fluorescently labeled CD80 or CD86, with or without high-concentration CTLA4-Ig (50 μg/ml) (fig. S3B). High CTLA4-Ig concentrations reduced cluster signals to background levels, indicating that ligands shifted from CD28 to CTLA4-Ig interactions (fig. S3B, two graphs on the left). Anti-CD28 blocking antibodies confirmed that cluster formation specifically depends on CD28 interactions (fig. S3B, two graphs on the right), validating our assay’s ability to distinguish CD28-mediated clustering from CTLA4-mediated diffuse signals.

**Fig. 2.**
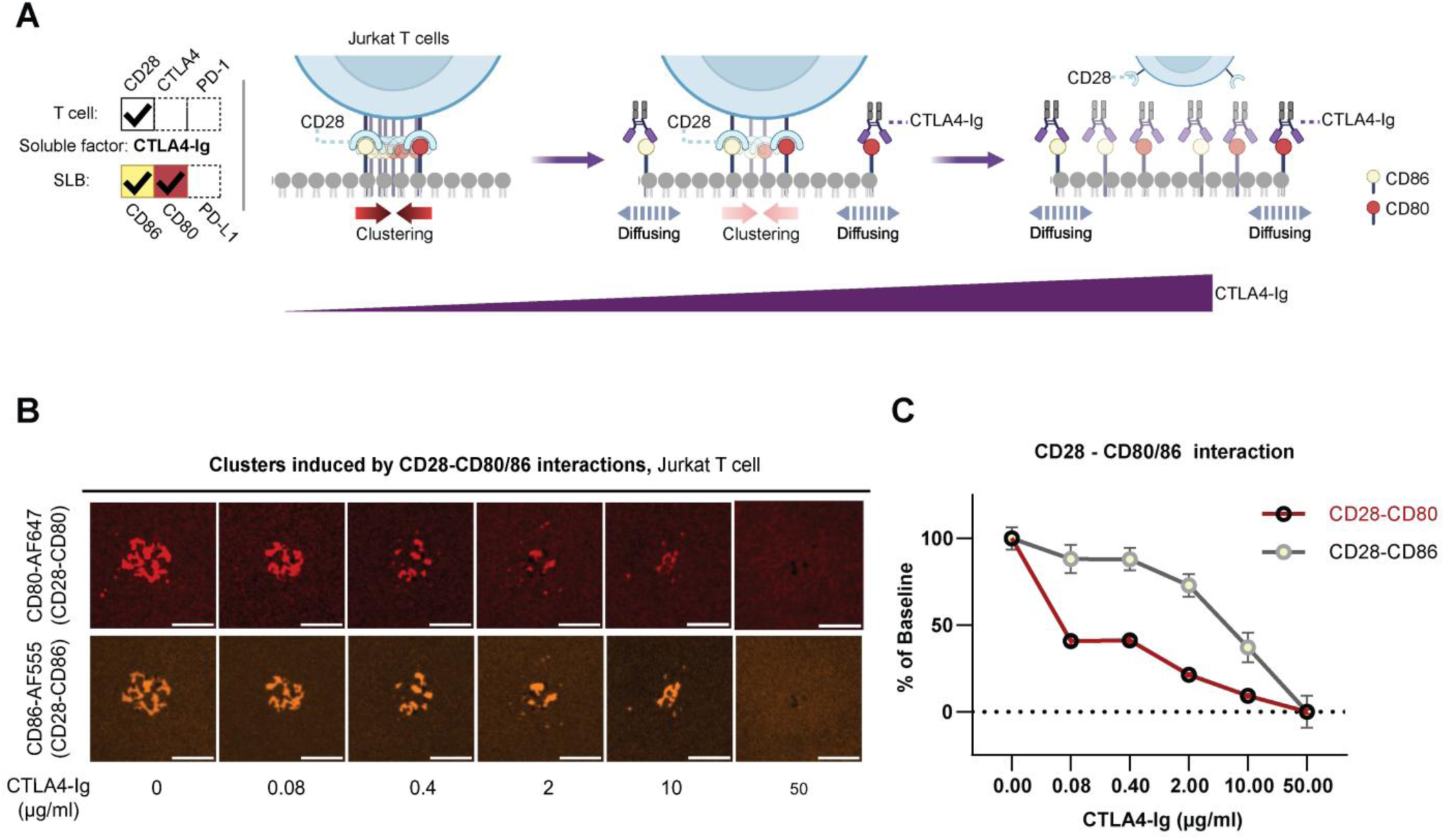
CTLA4 leads to increased interactions between CD28 and CD86 due to its higher affinity to CD80. **(A)** Schematic of soluble CTLA4-Ig competition assay with Jurkat T cells visualizing CD28-ligand interactions on SLBs containing anti-TCR VHH, ICAM-1, CD80, and CD86. **(B)** Representative images showing changes in CD28-CD80 and CD28-CD86 accumulation upon CTLA4-Ig addition. Scale bars: 5 µm. **(C)** Quantification of cluster intensities normalized to baseline (without CTLA4-Ig). Data is presented as means ± SEM (N_Cells_ >150, from two independent experiments).

We then incorporated both CD86 and CD80 on the lipid bilayer and incubated Jurkat T cells with varying concentrations of soluble CTLA4-Ig (Fig. 2A). We found that soluble CTLA4-Ig more readily competes with CD28 for binding to CD80 than CD86. Notably, even at a low CTLA4-Ig concentration of 0.08 μg/ml, CD80 cluster signals were reduced to approximately 50% of their initial intensity (Fig. 2B and C, ‘CD28-CD80’). This substantial reduction at a low concentration of CTLA4-Ig underscores the high affinity of CTLA4 for CD80 (*11*). As we increased the CTLA4-Ig concentration further, the decrease in CD80 clusters continued, ultimately reaching background levels at the highest concentration of 50 μg/ml. In contrast, CTLA4-Ig had a lesser impact on CD86-AF555 clusters. CD28-CD86 interactions remained less affected up to a CTLA4-Ig concentration of 0.4 μg/ml, where CD80 interactions were significantly impacted (Fig. 2B and C, ‘CD28-CD86’). These results demonstrate how a hierarchy of affinities promote ligand sharing between CD28 and CTLA4 through competitive interactions.

### Anti-PD-1 and anti-CTLA4 antibodies block ligand interactions to their receptors, with an unexpected loss of CD86 clusters with PD-1 blockade

Following our quantitative analysis of CTLA4’s effect on its ligands and CD28 interactions (Fig. 2), we expanded our investigation to include PD-1 and its ligand in the immune synapse (Fig. 3). We examined the interplay of CD80, CD86, and PD-L1 at the immunological synapse using stimulated primary CD3^+^ T cells from healthy donors, which expressed higher levels of checkpoint receptors PD-1 and CTLA4 compared to unstimulated T cells (fig. S4A), as expected (*22, 26*). We examined two different relative abundances of fluorescently labeled CD80, CD86 and PD-L1 in the SLB, a density of 200:200:200 /µm^2^ (1:1:1 CD80:CD86:PD-L1 ratio) and a density of 33:167:200 /µm^2^ (1:5:3 CD80:CD86:PD-L1 ratio) to reflect physiological relative abundances of these ligands on human antigen presenting cell (APC) (*52*).

**Fig. 3.**
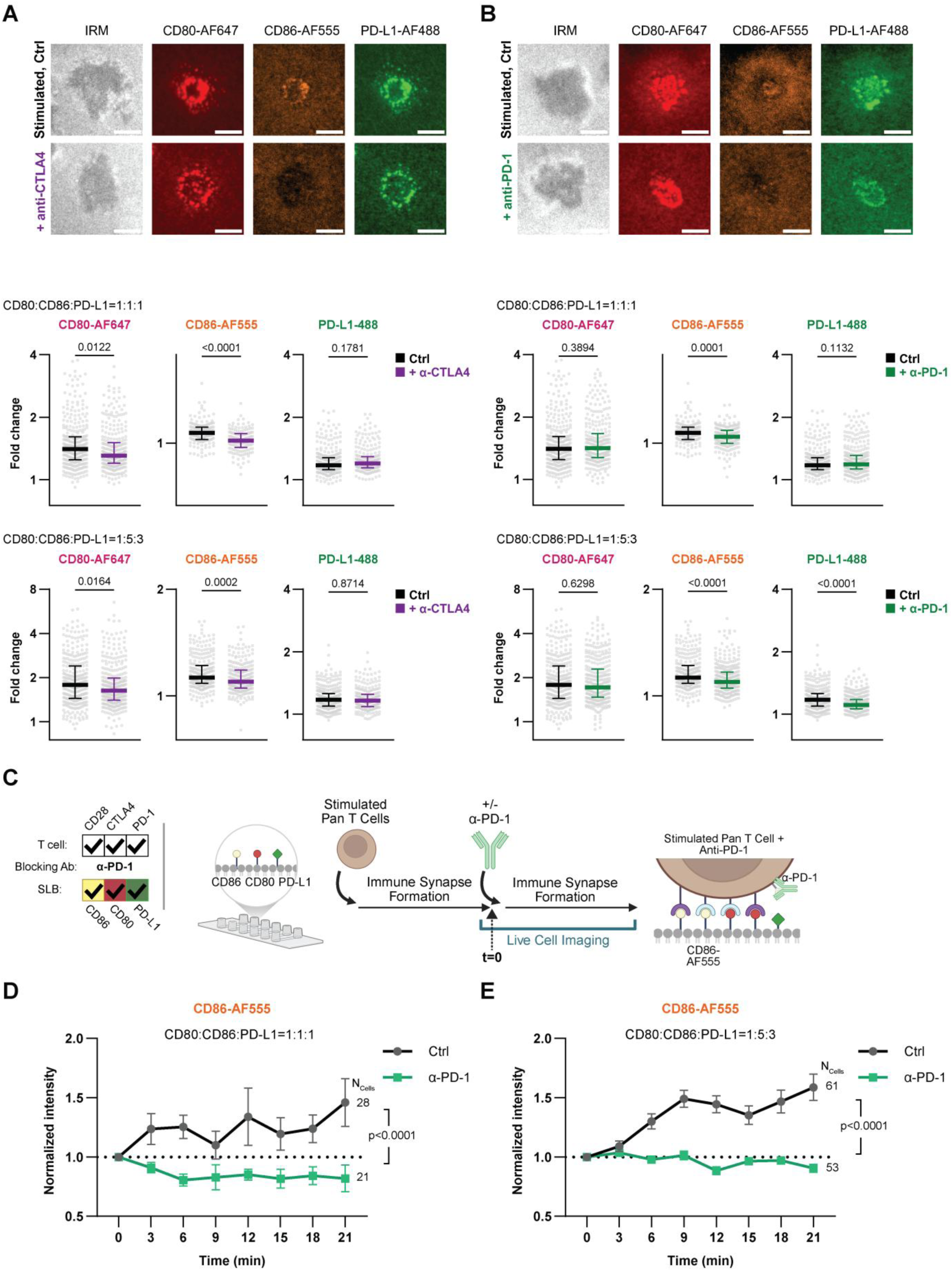
Anti-PD-1 and anti-CTLA4 antibodies block ligand interactions to their receptors, with an unexpected loss of CD86 clusters with PD-1 blockade. **(A and B)** Representative images of stimulated human primary CD3^+^ T cells forming immune synapses on SLBs containing anti-TCR VHH, ICAM-1, CD80-AF647, CD86-AF555, and PD-L1-AF488. From left to right are interference reflection microscopy (IRM) image and accumulation of CD80-AF647, CD86-AF555 and PD-L1-AF488 respectively - / + CTLA4 blockade (**A**) and - / + PD-1 blockade in (**B**). All representative images are from 1:1:1 ratio, Scale bars: 5 µm. Lower panels show quantification of CD80, CD86 and PD-L1 accumulation per synapse as a fold change over area of bilayer with no cell. The effects of CTLA4 blockade (**A**) and PD-1 blockade (**B**) on the accumulation of CD80, CD86 and PD-L1 were quantified at two different relative proportions of the ligands, CD80: CD86: PD-L1 = 1:1:1 and 1:5:3. Data presented as a dot column plot with median ± interquartile range (N_Cells_ > 200 from three independent experiments). P values determined by one-way ANOVA with multiple comparisons using Dunnett’s correction. **(C-E)** Real-time, live cell imaging to visualize the effects of PD-1 blockade on CD86 accumulation. **(C)** Schematic of live cell imaging setup. Stimulated T cells are incubated on SLBs containing anti-TCR VHH, ICAM-1, CD80, CD86-AF555, and PD-L1 for 24 minutes after which the anti-PD-1 antibody is added to the chamber. The cells are imaged for 21 minutes after antibody injection. (**D-E**) CD86 cluster intensity was quantified and normalized to the time point of antibody addition, depicted in the graph as t=0. The experiment was done under two different relative proportions of the ligands, CD80: CD86: PD-L1 = 1:1:1 (**D**) and 1:5:3 (**E**). Data is presented as means ± SEM (N_Cells_ shown in graphs). P values determined by two-way ANOVA.

We first examined the effects of CTLA4 blockade on ligand clustering using stimulated T cells (Fig. 3A and fig. S4B-C). In bilayers that contained either CD80 and PD-L1 or CD86 and PD-L1 in 1:1 ratios, we observed decreases in CD80 and CD86 clustering upon CTLA4 blockade (fig. S4B and C, ‘α-CTLA4’). We found a similar decrease in both CD80 and CD86 clustering upon CTLA4 blockade when all three ligands were present in the lipid bilayer at different proportions (Fig. 3A, ‘CD80-AF647’ and ‘CD86-AF555’). As expected, anti-CTLA4 blockade did not significantly influence PD-L1 clustering intensity (Fig. 3A, ‘PD-L1-AF488’). The clustering pattern of CD80 and PD-L1 is consistent with the localization of CD28 and PD-1 in peripheral microclusters and accumulation of CTLA4 in the cSMAC.

Next, we investigated the effects of PD-1 blockade on the accumulation of PD-L1, CD80 and CD86. Using primary activated T cells, we first compared the impact of PD-1 blockade when these cells were incubated with bilayers that contained CD86 and PD-L1 or CD80 and PD-L1 in 1:1 ratios (fig. S4B and C, ‘α-PD-1’). The amount of PD-L1 clustering observed on bilayers that contained CD80 and PD-L1 was much greater than on bilayers that contained CD86 and PD-L1 (fig. S4B and C, ‘PD-L1-AF488’). Upon PD-1 blockade, PD-L1 clustering reduced significantly on bilayers that contained CD86 (fig. S4B, ‘α-PD-1’), but there was no change in PD-L1 clustering in bilayers that contained CD80 (fig. S4C, ‘α-PD-1’). These experiments, show that the enhanced PD-L1 clustering observed in bilayers containing CD80 is due to the PD-L1-CD80 *cis*-interactions. There was limited impact of PD-1 blockade on PD-L1 clustering perhaps because released PD-L1 further interacted with CD80. CD80 and PD-1 compete for the same binding site on PD-L1 (*43, 53*). It is challenging to predict the relative occupancy of CD80 and PD-1 by PD-L1 because the two-dimensional affinities of PD-1 and CD80 for PD-L1 are not known.

We then investigated the impact of PD-1 blockade when all three ligands (CD80, CD86 and PD-L1) were present in the lipid bilayer using primary activated T cells (Fig. 3B). Upon PD-1 blockade, PD-L1 clustering was decreased when the ratio of CD80: PD-L1 was 1:3 but was unaffected when the ratio was 1:1 (Fig. 3B, ‘PD-L1-AF488’). This difference may reflect the influence of CD80-PD-L1 *cis*-interactions; While PD-1 blockade disrupts PD-1-PD-L1 interactions, the concurrent formation of CD28-CD80-PD-L1 complexes through CD80-PD-L1 *cis*-interactions can compensate for loss in PD-L1 clustering. CD80 clustering remained largely unchanged by anti-PD-1 treatment (Fig. 3B, ‘CD80-AF647’).

We, however, observed an unexpected decrease in CD86 clusters by PD-1 blockade in both conditions, where the ratio of the ligands (CD80: CD86: PD-L1) was 1:1:1 and 1:5:3 (Fig. 3B, ‘CD86-AF555’). To further validate our findings, we performed live cell imaging to visualize the impact of PD-1 blockade on CD86 clustering (Fig. 3C). CD86 clustering by primary stimulated T cells from healthy donors was monitored at two different CD80: CD86: PD-L1 ratios on the SLB. Our real time observations showed distinct patterns of CD86 accumulation between control and anti-PD-1 treated groups (Fig. 3D and E). In control conditions, CD86-AF555 intensity from each cell gradually increased from t=0, reflecting natural immune synapse formation and stabilization. In contrast, PD-1 blockade treated T cells showed minimal CD86 clustering increases over the same period, with some instances showing slight decreases. We validated our observations using a more physiological CD80:CD86 ratio (Fig. 3E). This ratio yielded results consistent with our initial observations, showing that anti-PD-1 blockade led to relatively decreased CD86 clustering compared to controls, where CD86 clusters continued to increase over time. These real-time imaging results (Fig. 3C-E) validate our single time point observations (Fig. 3B), confirming that PD-1 blockade reduces CD86 clustering over time.

Our hypothesis to explain these results is that upon PD-1 blockade, multiple interactions are re-organized: 1) the released PD-L1 forms more CD80-PD-L1 *cis*-heterodimers than CD80 homodimers, 2) these CD80-PD-L1 *cis*-heterodimers outcompete CD86 for binding to CD28, 3) the resulting decrease in CD80 homodimers lead to reduced binding to CTLA4. As shown in Fig 1, the relative binding of CD28 to CD80 and CD86 is unaffected by PD-L1. Therefore, CD80 released from CTLA4 plays an important role in the out competition of CD86 upon PD-1 blockade. To further explore this, we performed experiments in cells devoid of PD-1 and determined the relative ability of CD80 and CD86 to bind to CTLA4 and CD28 when PD-L1 is present.

### Binding of CD80 and CD86 to CTLA4 and CD28 in the absence of PD-1 but presence of PD-L1

Jurkat T cells express CD28 but are devoid of CTLA4 and PD-1. We generated a CTLA4 over-expressing Jurkat T cell line (CTLA4+ Jurkat) (fig. S5) and examined the influence of PD-L1 on CD86 clustering in the presence of CD80. We found that the extent of CD86 clustering decreased in the presence of PD-L1 (Fig. 4A and B) supporting our previous observation that CD80-PD-L1 *cis*-heterodimers induced by PD-1 blockade can outcompete CD86 for CD28 binding in the presence of CTLA4 (Fig. 3). However, these experiments did not directly show that CTLA4 binding to CD80 is reduced in the presence of PD-L1.

**Fig. 4.**
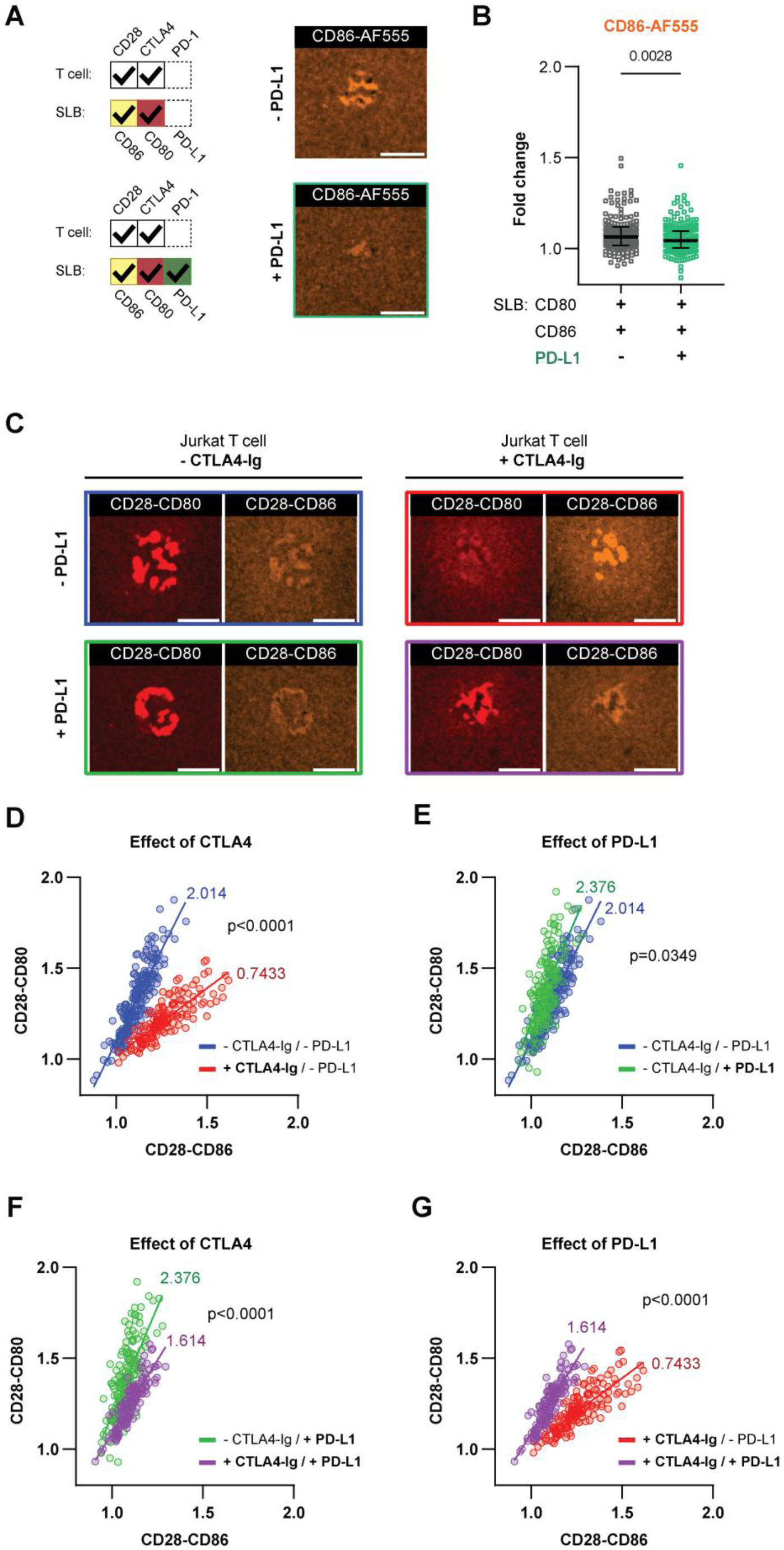
Binding of CD80 and CD86 to CTLA4 and CD28 in the absence of PD-1 but presence of PD-L1. **(A-B)** Changes in CD86 accumulation at immune synapses of CTLA4+ Jurkat T cells upon PD-L1 addition. **(A)** Schematic and representative images of CD86-AF555 accumulation of CTLA4+ Jurkat T cells forming immune synapses on SLBs containing anti-TCR VHH, ICAM-1, CD80, CD86-AF555, without (upper) or with (lower) PD-L1. Scale bars: 5 µm. **(B)** Quantification of CD86 cluster intensity at individual immune synapses without or with PD-L1 shown as a fold change over area of bilayer with no cells. Data presented as a column dot plot with medians ± interquartile range (N_Cells_=170 and 216, from three independent experiments). P value determined by unpaired t-test. **(C-G)** Soluble CTLA4-Ig assay visualizing changes in CD28-CD80 and CD28-CD86 interactions at immune synapses between Jurkat T cells and SLBs containing anti-TCR VHH, ICAM-1, CD80, CD86, with or without PD-L1. **(C)** Representative images showing CD28-CD80 and CD28-CD86 interactions without (left) or with (right) CTLA4-Ig, and without (upper) or with (lower) PD-L1. Scale bars: 5 µm. **(D-G)** Scatter plots of CD28-CD86 (x-axis) and CD28-CD80 (y-axis) fold accumulation, showing effects of CTLA4-Ig where PD-L1 is absent (**D**) or present (**F**) and showing the effect of PD-L1 where CTLA4 is absent (**E**) or present (**G**). Linear regressions were performed and values of slopes shown. P values determined by Analysis of Covariance (ANCOVA), comparing slope difference significance.

To investigate this, we performed experiments with Jurkat T cells that only express CD28 and examined the ability of soluble CTLA4-Ig to outcompete ligands from CD28 in the presence or absence of PD-L1 (Fig. 4C-G). This soluble CTLA4-Ig assay, characterized earlier in fig. S3B, ensures only CD28-mediated interactions would be visualized as clusters. Compared to the control group (Fig. 4C, ‘-CTLA4-Ig /-PD-L1’), CTLA4-Ig addition significantly enhanced CD86-AF555 fluorescent signals (Fig. 4C, ‘+CTLA4-Ig /-PD-L1’). However, when PD-L1 was additionally present on the SLB, CD80-AF647 fluorescent signals became relatively predominant (Fig. 4C, ‘+CTLA4-Ig /+PD-L1’). A scatter plot was generated using fold change values for CD80 (y-axis) and CD86 (x-axis) per cell. A slope greater than 1 suggests that majority of CD28 is bound to CD80 as seen in blue scatter plots (Fig. 4D); addition of CTLA4-Ig shifts this slope to less than 1 implicating that CTLA4-Ig competed away CD80 from CD28 and now more of CD28 is bound to CD86 (Fig. 4D). In the absence of CTLA4-Ig, PD-L1 only marginally increases binding of CD28 to CD80 (Fig. 4E). In the presence of PD-L1, CTLA4-Ig had a low impact on the preference of CD28 binding to CD80 (Fig. 4F), in contrast to the significant impact of CTLA4-Ig seen in the absence of PD-L1 (Fig. 4D). In the presence of CTLA4-Ig, the slope was highly sensitive to the presence of PD-L1, changing from greater than 1 (+PD-L1) to less than 1 (-PD-L1) (Fig. 4G). This indicates that CD80-PD-L1 *cis*-heterodimers diminish CD80’s capacity to bind CTLA4. Our orthogonal validation through directly adding PD-L1 to PD-1-absent systems shows that CD80-PD-L1 heterodimer reduces CD28-CD86 interactions while increasing CD28-CD80 interactions, but only in the presence of CTLA4. This demonstrates that CTLA4-CD80 disruption can be triggered by enhanced CD80-PD-L1 *cis*-interactions.

### PD-1 enrichment at the immune synapse as a surrogate for engaged PD-1 reveals the impact of CD80-PD-L1 *cis*-interactions

Studies have shown that PD-1 localizes to TCR microclusters only when it is bound to PD-L1 (*30, 54*). In previous experiments (Fig. 3), interpreting PD-L1 cluster intensity changes was challenging because PD-L1-AF488 signals could originate from either PD-1-PD-L1 interactions or CD28-CD80-PD-L1 ternary complexes. We employed a PD-1-binding scFv-AF647 reagent that does not compete with our PD-1 blocking antibody (*55*), enabling direct detection of PD-1 accumulation to immune synapses using TIRFM imaging (Fig. 5). Using primary activated T cells, we investigated the impact of CD80 on the accumulation of PD-1 in the immune synapse by preparing bilayers that contained CD86 and PD-L1-AF488 in addition to anti-TCR VHH and ICAM-1. Additionally, we investigated the impact of PD-1 and CD28 blockade. All experiments were performed at two different relative abundances mirroring experiments performed in Fig.3. We first observed that the amount of PD-1 in the synapse was lower in the presence of CD80, likely due to a lower proportion of PD-1 bound to PD-L1 (Fig. 5A (i) and 5B (i)) consistent with the shared binding site on PD-L1 for PD-1 and CD80. Upon PD-1 blockade, the amount of PD-1 detected in the immune synapse decreased in the presence and absence of CD80 (Fig. 5A (ii) and 5B (ii)). Of note that the PD-L1-AF488 clustering showed only modest reduction, with substantial amount of PD-L1 clustering remaining at the immunological synapse in previous experiments (Fig. 3B, ‘CD80: CD86: PD-L1=1:1:1’) and in this one (fig. S6). CD28 blockade, which may have potentially generated more available CD80 did not have any impact PD-1 intensity in the immune synapse (Fig. 5A (iii) and 5B (iii)). The dual blockade of CD28 and PD-1 did not have any impact on PD-1 intensity in the immune synapse (Fig. 5A (iv) and 5B (iv)) but dramatically reduced PD-L1 accumulation (fig. S6, ‘Combo’). These results directly demonstrate that CD86 reduction in Fig. 3 results from increased CD28-CD80-PD-L1 ternary complex formation following PD-1-PD-L1 disruption.

**Fig. 5.**
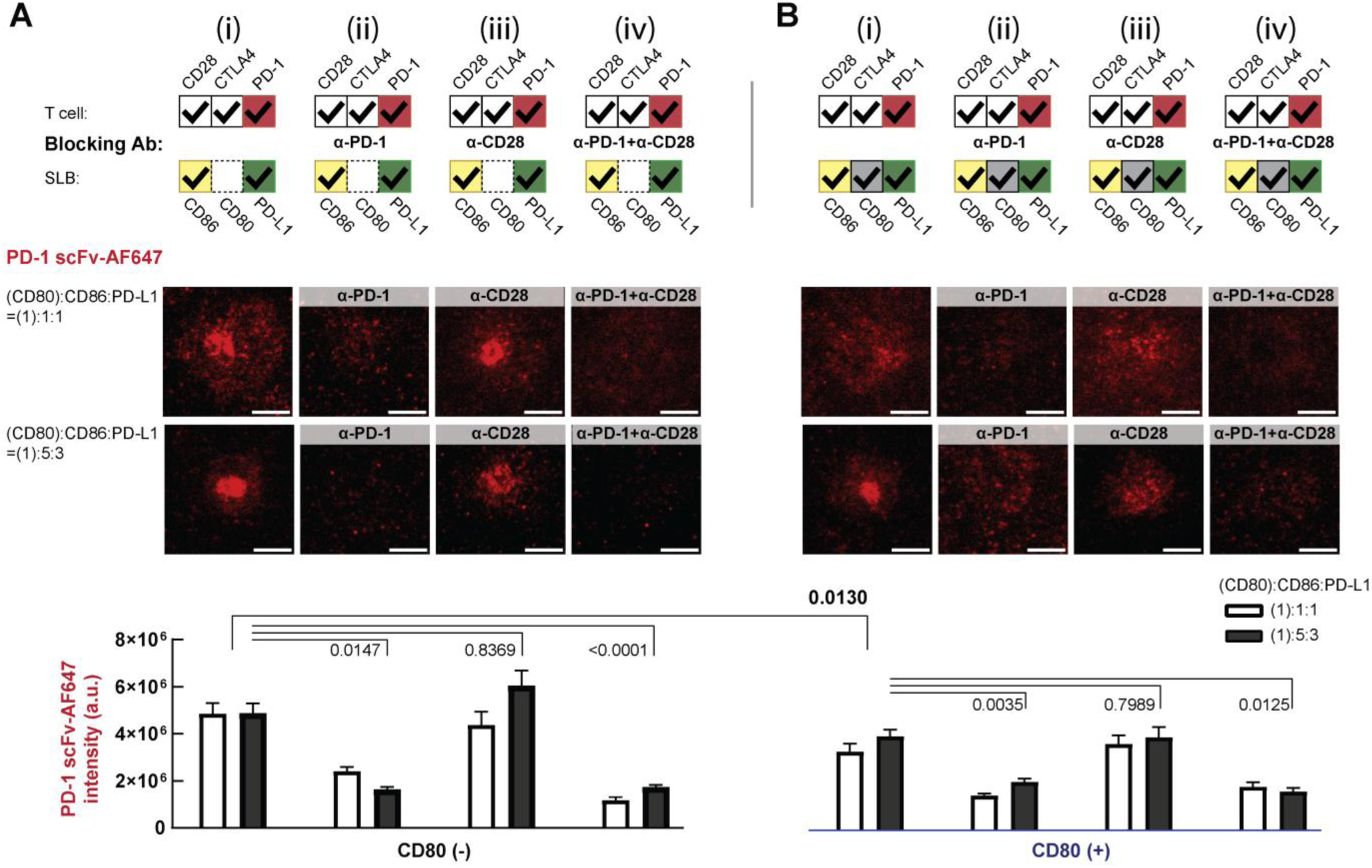
PD-1 enrichment at the immune synapse as a surrogate for engaged PD-1 reveals the impact of CD80-PD-L1 *cis*-interactions. **(A-B)** Stimulated primary CD3^+^ T cells forming immune synapses on SLBs containing anti-TCR VHH, ICAM-1, and two ratios of CD86, PD-L1-AF488, without CD80 (**A**) or with CD80 (**B**). Cells were imaged using total internal reflection fluorescence microscopy (TIRFM) in the presence of non-blocking and non-competing AF647-labeled anti-PD-1 scFv to visualize PD-1. Representative images showing the polarization of PD-1 (PD-1 scFv-AF647) at immune synapses and within microclusters (**A**(i) and **B**(i)). The effects of PD-1 blockade (**A**(ii) and **B**(ii)), CD28 blockade (**A**(iii) and **B**(iii)), and dual blockade of CD28 and PD-1 (**A**(iv) and **B**(iv)) were also investigated. Scale bars: 5 µm. Averaged cluster intensities for PD-1 were determined for each condition. Data presented as means ± SEM for two ligand ratio compositions (N_Cells_ >120 from three independent experiments per condition). P values from experiments with (CD80):CD86: PD-L1=(1):1:1, determined by one-way ANOVA with multiple comparisons. See also fig. S6 for the quantification of PD-L1 accumulation.

### The relative costimulatory potential of CD80 and CD86

Given the weaker affinity of CD86 for CD28, previous studies have suggested that CD28-CD86 interactions generate weaker signals compared to CD28-CD80 interactions (*11*). If this differential signaling capacity holds true at immune synapses, then PD-1 blockade that shifts the CD28-ligand balance from CD86 toward CD80 (Fig. 3-5) should enhance T cell activation signaling beyond the effects of blocking PD-1 inhibitory signaling. To investigate whether CD80 promotes more efficient immunological synapse formation than CD86 at the same ligand densities on APCs, we employed interference reflection microscopy (IRM) coupled with automated segmentation analysis to quantify the quality of cell adhesion by primary human T cells on SLBs over time. SLBs contained anti-TCR VHH-His, ICAM-1^ECD^-His, and either CD80^ECD^-His or CD86^ECD^-His (Fig. 6A-C).

**Fig. 6.**
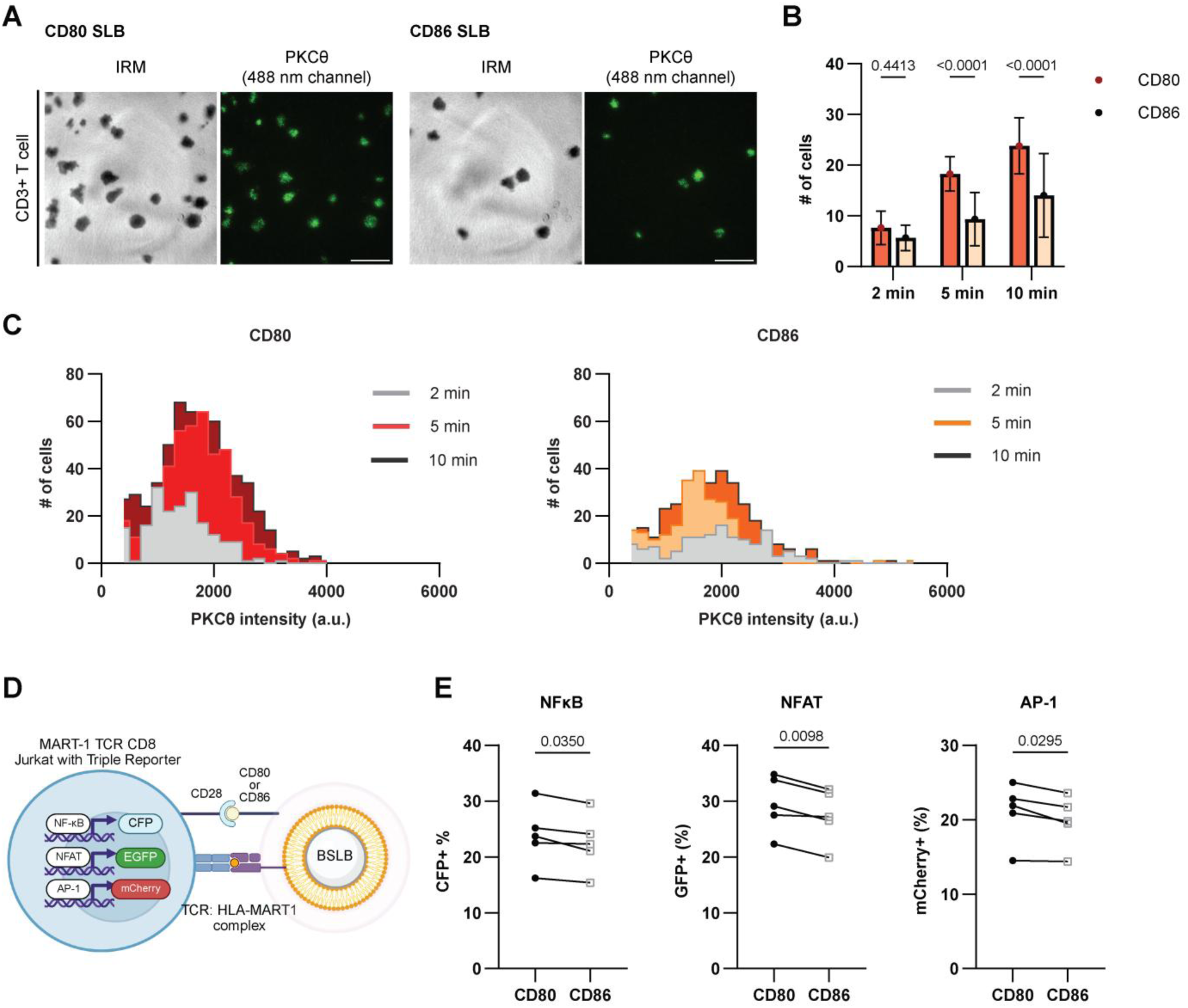
The relative costimulatory potential of CD80 and CD86. **(A-C)** Comparison of T cell adhesion efficiency on CD80 versus CD86 SLBs. **(A)** Representative images of freshly isolated human primary CD3^+^ T cells forming immune synapses on SLBs containing anti-TCR VHH, ICAM-1, ∼20 /µm^2^ density of CD80 (left) or CD86 (right), 5 minutes post-injection. Scale bars: 25 µm. **(B)** Quantification of successfully attached T cells per field of view using automatic segmentation based on IRM images. Data presented as bar plots with mean ± S.D. (N=24 images from three independent experiments). P values determined by unpaired t-test comparing CD80 and CD86 SLBs at each time point. (**C**) Histogram showing the frequency distribution of PKCθ average intensity measurements across individual immune synapses on CD80 or CD86 SLBs at each time point. Data pooled from three independent experiments. **(D)** Schematic of T cell activation assay using antigen-specific triple-reporter Jurkat T cell line (see Methods for more details) and bead-supported lipid bilayers (BSLBs) containing ∼200 /µm^2^ density of CD80 or CD86. **(E)** Flow cytometry analysis of activation of triple reporter Jurkat T cell showing the impact of CD80 vs CD86 co-stimulation, N=5 independent experiments. P values determined by paired t-test.

We fixed T cells on the SLBs containing either CD80 or CD86 at 0, 2, 5, and 10-minute time points and quantified the number of cells achieving high-quality adhesion per field based on IRM pattern. Notably, at 5 and 10-minute time points, significantly more cells formed immunological synapses on CD80 SLBs compared to CD86 SLBs (Fig. 6A and B). Signals downstream of CD28 lead to the recruitment of PKCθ to the immune synapse. While there was a difference observed in the fraction of cells adhering to bilayers containing CD80 or CD86, there was no difference in the amount of PKCθ clustering at each immunological synapse (Fig. 6C). These results suggest that during the initial stages of T cell-APC contact, CD28 clustering and signaling through CD80 is more efficient than through CD86 leading to better T cell APC adhesion. Note that the freshly isolated T cells used here expressed CD28 but not CTLA4.

After confirming that CD80 promotes more efficient T cell adhesion compared to CD86 during initial CD28 interaction, we sought to investigate longer-term T cell-APC interactions. We generated bead-supported lipid bilayers (BSLBs) coated with human leukocyte antigen A2 molecule (HLA-A2) coupled with melanoma antigen recognized by T-cells 1 (MART-1) along with MART-1-recognizing TCR-transduced CD8^+^ Jurkat T cells to induce antigen-specific Jurkat-BSLB interactions (Fig. 6D). This BSLBs approach was essential for our mechanistic studies as it allows precise control over ligand composition and density and enabled accurate comparison of CD80 versus CD86 effects on T cell activation without interference from other co-stimulatory or co-inhibitory molecules. The MART-1 TCR-T Jurkat T cells used here also expressed fluorescent reporters for downstream transcription factor activity following T cell activation (NFκB-CFP, NFAT-GFP, and AP-1-mCherry) (*56, 57*), whose expression can be monitored by flow cytometry. We confirmed that these T cells were activated in an antigen-specific manner (fig. S7). We prepared BSLB that contained MART-1-peptide-HLA-A2 complexes, ICAM-1, and either CD86 or CD80. After 18-hours of co-culture we measured the activity of NFκB, NFAT, and AP-1 by flow cytometry. Consistent with our adhesion data we found that CD80 modestly induced better activity of NFκB, NFAT, and AP-1 compared to CD86 (Fig. 6E).

### PD-1 blockade enhances T cell activation through two distinct mechanisms

Having established a system where we could measure the relative functional effects of CD80 and CD86, we sought to use it to test the hypothesis that PD-1 blockade has a dual impact on T cell activation, one through blocking PD-1 inhibitory signals and second through enhancement of CD28-CD80 interactions by disrupting CTLA4-CD80 complexes through increased CD80-PD-L1 *cis*-interactions. To test this, we performed co-culture experiments using CTLA4^+^ PD-1^+^ MART-1 TCR-transduced Jurkat T cells (fig. S8A) with BSLBs coated with HLA-MART-1 complexes and ligands (Fig. 7A). If PD-1 blockade solely functions by disrupting PD-1-PD-L1 complexes and eliminating PD-1 signaling with no contribution from CD80-PD-L1 *cis*-interactions then T cell activation should be equivalent between conditions with CD80 alone versus CD80/PD-L1 with anti-PD-1 antibody (Fig. 7A), since both conditions would lack PD-1 inhibitory signaling. As a control we compared T cell activation by BSLB that contained CD86 and CD86+PD-L1 with PD-1 blockade.

**Fig. 7.**
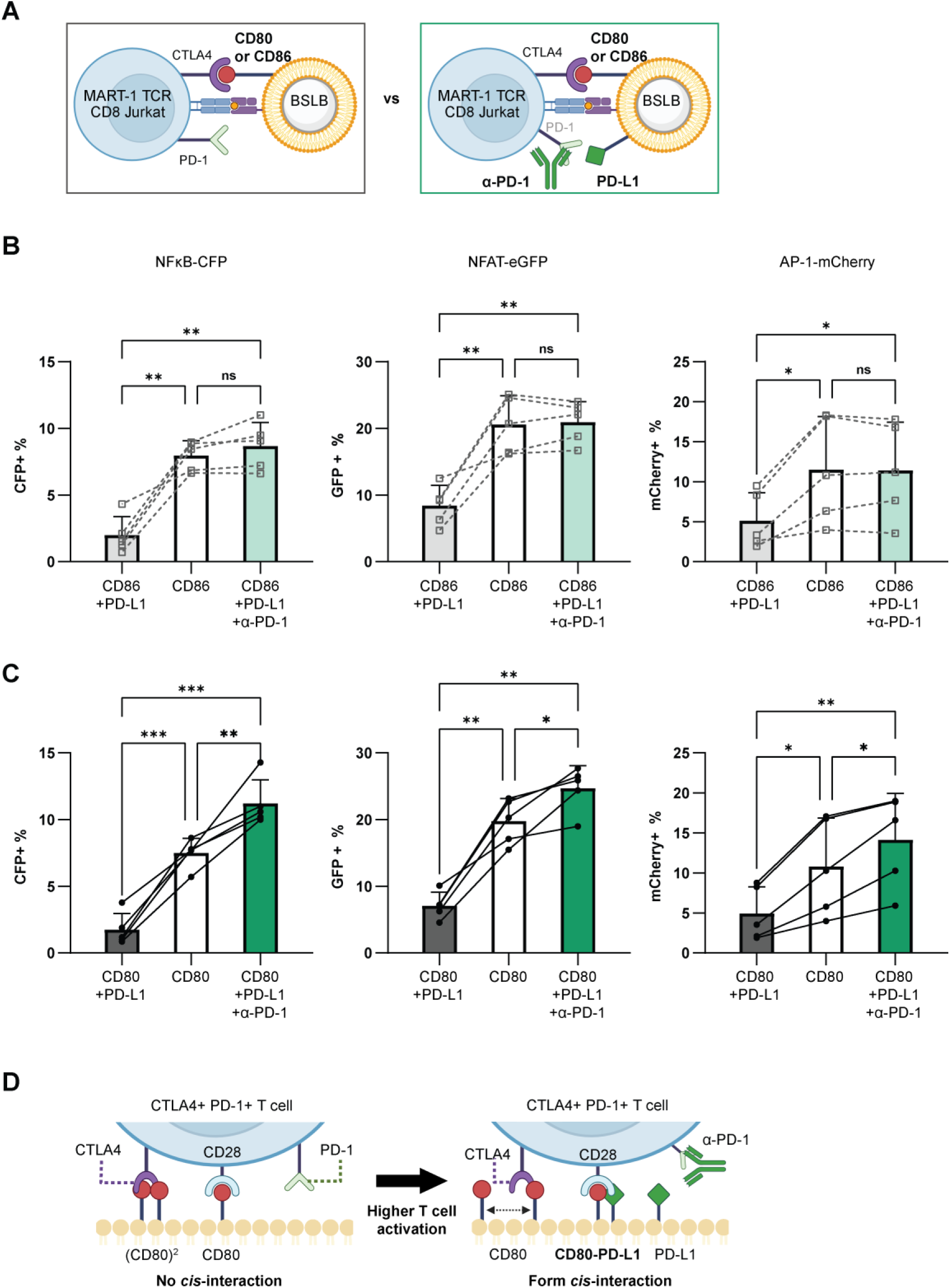
PD-1 blockade enhances T cell activation through two distinct mechanisms. **(A)** Schematic of co-culture assay using antigen-specific triple-reporter Jurkat T cell line and BSLBs. BSLBs are coated with HLA-MART1 complexes, CD80 or CD86, without PD-L1 (left) and with PD-L1 +/- PD-1 blockade (right) (CD28 not shown for simplicity). **(B-C)** Flow cytometry analysis of T cell activation marker expression under different BSLB compositions and antibody treatments. **(B)** Experiment comparing activation of triple reporter Jurkat cells in response to CD86 versus CD86+PD-L1+anti-PD-1 conditions (N = 5 independent experiments). P values determined by paired t-test. **(C)** Similar experiment as in B but comparing activation in response to CD80 versus CD80+PD-L1+anti-PD-1 conditions (N = 5 independent experiments). P values determined by paired t-test. **(D)** Schematic representation of proposed new model based on experimental findings.

We first investigated T cell activation by the CTLA4^+^ PD-1^+^ MART-1 TCR-transduced Jurkat T cells with BSLBs coated with HLA-MART-1 complexes, ICAM-1 and CD86 with or without PD-L1 (Fig. 7B and fig. S8B). PD-L1 addition significantly reduced T cell activation as measured by expression of reporters for NF-kB, AP-1 and NFAT highlighting the inhibitory effect of PD-1 signaling (Fig. 7B, ‘CD86+PD-L1’), PD-1 blockade in CD86+PD-L1 condition restored T cell activation to the same level as CD86 alone (Fig. 7B, ‘CD86+PD-L1+α-PD-1’). We then investigated BSLBs that contained CD80 or CD80+PD-L1 (Fig. 7C and fig. S8C). As seen with the CD86 conditions, the addition of PD-L1 to CD80 containing BSLB significantly reduced T cell activation (Fig. 7C, ‘CD80+PD-L1’). PD-1 blockade in the CD80+PD-L1 condition showed significantly higher T cell activation than in the CD80 alone condition (Fig. 7C, ‘CD80+PD-L1+α-PD-1’). These results support our hypothesis that PD-1 blockade enhances T cell activation through two distinct mechanisms: one through blockade of inhibitory signaling and second through increased formation of CD80–PD-L1 *cis*-heterodimers that have increased binding to CD28 than CTLA4 (Fig. 7D).

## Discussion

The plethora of co-stimulatory and co-inhibitory pathways have a profound impact on regulation of immune responses. These pathways are difficult to study as most proteins in these pathways are membrane proteins and tools to determine two-dimensional affinities are limited and subject of intense debate (*58*). In addition, both *cis* and *trans* interactions among pathway members contribute to activity (*46, 59, 60*). For these reasons, a predictive and quantitative understanding of how CD28, CTLA4, and PD-1 receptors compete for their ligands in physiological conditions remains limited. While individual receptor-ligand interactions have been extensively characterized (*3, 5, 15, 61*), the critical challenge lies in determining the individual receptor occupancy when multiple ligands are simultaneously engaged. The *cis*-interaction between CD80 and PD-L1 further adds to the challenge of accurately determining the receptor occupancy for these receptors (*45*). Antibody mediated blockades of PD-1 and CTLA4 can have different effects in priming and effector phases of T cells because the relative expression of CD86, CD80 and PD-L1 in the two contexts could be very different. During early phase of priming, T cells may encounter APCs that have excess of CD86 relative to CD80 and PD-L1 (*2, 62*) while during later phases of priming, APCs may have upregulated CD80 and PD-L1 (*2, 5*). During the effector phase, T cells may encounter cells that are very low in CD80 and CD86, such as virus infected cells or tumor cells (*63, 64*). The contribution of the CD28, CTLA4 and PD-1 pathway to T cell activation will thus depend on several factors: the relative ability of CD86 and CD80 to deliver co-stimulatory signals, the competition between CTLA4 and CD28 for their ligands, the inhibitory signals delivered by PD-1–PD-L1 interactions and how CD80–PD-L1 *cis*-interactions regulate receptor occupancies. While these factors have been individually studied, this is the first investigation of them in an integrated study.

In this study, we visualized CD80, CD86, and PD-L1 interactions and quantified competitive interactions among them by imaging the T cell–artificial APC immune synapse formed between a T cell and a glass supported lipid bilayers. Consistent with known binding affinities (*11*), we confirmed that CD80 maintains dominant interactions with CD28, requiring approximately one order higher CD86 density to achieve comparable binding. Sansom and colleagues reported that when T_regs_ that express CD28 and high levels of CTLA4 interact with an APC that express both CD80 and CD86, CD28 is primarily occupied by CD86 because of the higher affinity of CTLA4 for CD80 and its preference for binding to dimers of CD80 (*13*). We were able to demonstrate the same phenomenon by measuring the ability of CTLA4-Ig to out compete CD80 or CD86 interactions from CD28 (Fig. 2). We found that very low concentrations of CTLA4-Ig could displace CD80 from CD28 while, much higher CTLA4-Ig concentrations were required to displace CD86 from CD28. We made an unexpected observation when we examined the role of PD-1 in this network. We found that PD-1 blockade led to reduced accumulation of CD86, when CD86 is not known to directly bind to PD-1. We hypothesized that this occurred through increased CD80-PD-L1 *cis*-interactions that led to the displacement of CD86 from CD28 (Fig. 3-4). We additionally showed that these CD80-PD-L1 heterodimers preferred to bind to CD28 over CTLA4 due to the preference of CTLA4 for CD80 dimers. While it has been shown that CTLA4 can disrupt CD80-PD-L1 *cis*-interactions through re-organizing CD80 on the cell surface (*65*), our results would argue that this would be limited by the equilibrium between CD80-PD-L1 heterodimers and CD80 homodimers as CTLA4 has reduced binding to CD80-PDL1 heterodimers. Using selective PD-1 detection, we directly visualized PD-L1 partner switching from PD-1 to CD80 upon PD-1 blockade, demonstrating that liberated PD-L1 molecules strengthen CD28-CD80-PD-L1 ternary complexes that outcompete CD28-CD86 interactions (Fig. 5). Functionally, we showed that PD-1 blockade operates through two mechanisms: enhancing CD28-CD80 co-stimulation through redistribution of ligands and conventional PD-1 signaling blockade (Fig. 7). Our bead-supported lipid bilayer systems provided the precise control over ligand composition and density while excluding confounding effects from other immune molecules, which was essential for dissecting the dual mechanisms of PD-1 antagonism.

Our study builds upon the work from Sugiura et al. and Zhao et al. who demonstrated that CD80 and PD-1 share the binding site on PD-L1 (*43, 44*). Zhao et al expressed CD80 or PD-L1 at very high levels to conclude that in the presence of very high levels of CD80, PD-1–PD-L1 interactions may not occur (*44*). Their studies did not include CD86 and using more physiological levels of CD80 and PD-L1 we are able to demonstrate that significant PD-1– PD-L1 interactions still exist (Fig. 5). We have shown that CD80–PD-L1 *cis*-interactions trigger a cascade of competitive effects upon PD-1 blockade; released PD-L1 forms more *cis*-interactions that have reduced capacity to bind to CTLA4 and increased capacity to bind to CD28, in effect demonstrating that PD-1 blockade also leads to partial CTLA4 antagonism explaining perhaps why PD-1 blockade has been so effective in the clinic. Functionally, we were able to show that TCR signaling upon PD-1 blockade of bilayers containing CD80 and PD-L1 was more than bilayers that contained just CD80 (Fig. 7). This likely reflects the reduced capacity of CD80–PD-L1 heterodimers to bind to CTLA4.

Our observations of rapid redistribution of ligands upon PD-1 blockade, need to be considered alongside slower regulatory processes such as CTLA4-mediated trans-endocytosis. CTLA4-mediated trans-endocytosis is reported to continuously remove CD80 from the surface of APCs over time, potentially shifting the balance toward CD86-dominant interactions (*47, 48, 66*). This slow membrane remodeling process contrasts with the minute-timescale ligand redistribution we observed upon PD-1 blockade, highlighting that checkpoint receptor regulation operates across multiple time scales. We speculate that moderate CD80 depletion might initially enhance the relevance of our mechanism by making CD28-CD86 interactions more prominent, thereby increasing the potential impact of PD-1 blockade-mediated competitive redistribution toward CD28-CD80 binding. However, if trans-endocytosis proceeds to near-complete CD80 removal or eliminates CTLA4-expressing cells entirely, our competitive mechanism would likely become irrelevant as the molecular players required for CD80-PD-L1 *cis*-interaction and CTLA4-mediated competition would be absent. How this dynamic interplay between rapid PD-1 blockade-mediated ligand redistribution and slower membrane remodeling intersects with our newly identified mechanism represents a critical area for future study, as it may determine the temporal window during which PD-1 blockade can effectively enhance CD28-CD80 co-stimulation.

Beyond these temporal considerations, the competitive ligand dynamics we identified have important implications for understanding how different APC types regulate T cell activation *in vivo*. Our quantitative analysis across varying CD80: CD86: PD-L1 ratios enabled precise determination of CD28 binding complex composition, revealing how both CTLA4 presence and CD80/PD-L1 availability serve as quantitative determinants that can reverse CD28’s preferential binding partners. These findings suggest that the distinct expression profiles of CD80, CD86, and PD-L1 on different APC subsets fundamentally determine the co-stimulatory signaling balance *in vivo*. Given that human dendritic cells, macrophages, and B cells exhibit distinct CD80:CD86 expression ratios that shift dramatically upon activation (*5, 67*), our findings predict that T cell responses will vary significantly depending on the specific APC type and activation state encountered in addition to the co-expression of other co-stimulatory and co-inhibitory molecules. This APC-dependent variability becomes particularly critical in tumor microenvironments, where malignant and stromal cells can express aberrant ligand ratios with additional complexity from intra- and inter- patient heterogeneity (*68*). In such contexts, PD-1 blockade efficacy depends critically on local ligand composition rather than PD-L1 expression alone, potentially explaining why PD-L1 remains an imperfect biomarker for anti-PD-1 therapy responses (*34, 69, 70*). Future computational modeling studies incorporating the quantitative parameters we identified will be essential for predicting optimal ligand ratio thresholds and determining which microenvironmental compositions favor the competitive enhancement mechanism.

In conclusion, our findings establish that PD-1 blockade operates through previously unrecognized competitive principles that extend far beyond simple pathway inhibition. By revealing that PD-1 antagonism operates through dual mechanisms, conventional inhibitory signal relief and competitive enhancement of CD28-CD80 co-stimulation, this work provides a new framework for understanding T cell activation in consideration of systemic network effects. The discovery that blocking one checkpoint pathway can paradoxically strengthen alternative receptor interactions opens new approaches for rational immunotherapy design and explains variable treatment responses observed clinically. Moving forward, optimizing checkpoint therapies will require network-level analysis that considers how interventions reshape the entire immune synapse rather than simply blocking individual molecular targets.

## Materials and Methods

### Human T cell isolation, activation and culture

Leukopaks from healthy adult donors were purchased from commercial suppliers (StemExpress, BioIVT, StemCell Technologies) with donor written informed consent and all materials were de-identified prior to receipt. Peripheral blood mononuclear cells (PBMCs) were isolated using ficoll gradient.

T cells were enriched from frozen human PBMCs using the Human Pan T Cell Enrichment Kit (StemCell Technologies, 19051) according to the manufacturer’s instructions. Activated T cells were generated by stimulating PBMCs with plate-bound anti-CD3 antibody (clone: OKT3, Biolegend). Plates were coated with 1 μg/ml of anti-CD3 antibody diluted in PBS and incubated overnight at 4°C. Human PBMCs were thawed, washed, and resuspended at a concentration of 2 × 10⁶ cells/ml in complete T cell culture medium (RPMI-1640 (Gibco 22400089) supplemented with 5% Human AB serum, 1% sodium pyruvate, and 1% penicillin/streptomycin). Cells were plated onto anti-CD3-coated 96-well plates and incubated for 72 hours at 37°C in a 5% CO₂ incubator. Following activation, CD3⁺ T cells were isolated using the Human Pan T Cell Enrichment Kit (StemCell Technologies, 19051). Parallel negative control experiments were performed using uncoated 96-well plates to assess differences between unstimulated and stimulated T cell populations.

### Cell lines and cultures

Jurkat T cells were obtained from the American Type Culture Collection (ATCC) (TIB-152) and were cultured in complete RPMI-1640 medium (RPMI-1640 medium (Gibco 22400089) supplemented with 10% fetal bovine serum (FBS) (Gibco), and 1% penicillin/streptomycin (Gibco)). CTLA4+ Jurkat T cells were generated by transfecting parental Jurkat T cells with a CTLA4-Blasticidin encoding lentivirus construct. CTLA4+ Jurkat T cells were maintained under antibiotic selection using 8 μg/ml blasticidin in complete RPMI-1640 medium.

MART-1 TCR CD8⁺ Jurkat T cells were engineered based on the TCR-negative Jurkat 76 cell line that was modified by lentiviral transduction to generate triple reporter-expressing cells (*71*). The expression constructs encoding human CD8 (alpha and beta) and MART-1 TCR (clone DMF4, against ELAGIGILTV) were virally transduced. These cells were subsequently transduced with PD-1 and CTLA4 expression constructs to create multi-transgenic cell lines. Engineered cell lines were maintained under antibiotic selection using 10 μg/ml blasticidin and 2 μg/ml puromycin in complete RPMI-1640 medium to ensure stable transgene expression.

All cell lines were maintained at 37°C in a humidified incubator with 5% (v/v) CO_2_. Parental cell lines (Jurkat T cells and Jurkat 76 Triple Reporter with MART-1 TCR/CD8) were periodically tested for mycoplasma contamination by cell services core team.

### Recombinant proteins

Soluble recombinant CTLA4-Ig (AcroBioSystems, CT4-H5255), monomeric biotinylated MART-1 peptide loaded HLA (MLB, TB-0009-M), and streptavidin (Thermo Fisher Scientific, S888) were obtained from commercial sources.

12×histidine-tagged recombinant proteins were produced by GenScript’s gene synthesis and protein purification services. These included extracellular domains (ECDs) of ICAM-1, CD80, CD80-KKCK (for site specific labeling, (*72*)), CD86, PD-L1, and anti-TCR VHH (clone 56G05) (*73*). Proteins were expressed by transfecting pcDNA3.4 sub-cloned plasmids into CHO cells and purified using HisTrap Fast Flow Crude Columns. Protein purity and mass were characterized by SEC-HPLC and LC-MS, respectively. Protein sequences are provided in Supplementary Information. Recombinant proteins were fluorescently labeled, except for functional assays, using amine reactive or cysteine reactive AlexaFluor labeling products (Thermo Fisher Scientific): AlexaFluor 488 (A10235), AlexaFluor 555 (A20174), AlexaFluor 647 (A20173). To minimize over-labeling of CD80, the CD80-KKCK protein was labeled with the cysteine reactive AlexaFluor 647 maleimide (A20347).

To visualize PD-1 a non-blocking anti-PD-1 scFv (clone 19, mAb12) (*55*) was produced by GenScript’s gene synthesis and protein purification services and labeled with AlexaFluor 647 labeling kit (Thermo Fisher Scientific). This scFv clone is also non-competing with the PD-1 blocking antibody (clone LO115) used here. This protein was expressed by transfecting sub-cloned pcDNA3.4 plasmid into CHO cells, purified using Anti-DYKDDDK G1 Affinity Resin, and characterized for purity by SDS-PAGE, SEC-HPLC, and CE-SDS. Protein sequence is provided in Table S1.

### Antibodies

Blocking antibodies including anti-CTLA4 (Tremelimumab), anti-PD-1 (LO115 bivalent), anti-PD-L1 (Durvalumab), and anti-CD28 monovalent (clone ANC28.1) (*74*) were produced in-house using the established plasmids for expression. Anti-CTLA4 (Tremelimumab) and anti-PD-1 (LO115 bivalent) were previously characterized (*75*). Characterization of anti-PD-L1 (Durvalumab, also known as MEDI4736) has also been previously reported (*76*). Briefly, encoding gene plasmids of human IgG1 triple mutant (TM, L234F/L235E/P331S) (*77*) mAb anti-CTLA4, anti-PD-1, and anti-PD-L1 antagonistic antibodies were transiently expressed in CHO-G22 cell lines and antibodies were purified using protein A column. Following buffer exchange and size exclusion chromatography was conducted. Protein purity and mass were characterized by SEC-HPLC and LC-MS, respectively.

The following anti-human antibody for flow cytometry was obtained from BD Biosciences: BV480 CD28 (CD28.2). The following anti-human antibodies were obtained from Biolegend: BV711 CD3 (SK7), BV605 isotype control (MOPC-21), BV605 CD25 (BC96), PE isotype control (MOPC-173), PE CTLA4 (BNI3), PerCP/Cy5.5 isotype control (MOPC-21), PerCP/Cy5.5 CD69 (FN50), APC isotype control (MOPC-21), APC PD-1 (EH12.2H7).

Anti-PKCθ primary antibody was purchased from Cell Signaling Technology (86952SF) and labeled with AlexaFluor 488 labeling kit (Thermo Fisher Scientific, A10235).

### Flow cytometry

For characterization of expression of receptors and activation in T cells, cells were resuspended at approximately 2 × 10⁶ cells/ml and washed in PBS. Cells were stained with Live/Dead Zombie NIR Fixable Viability Kit (BioLegend, 423105) according to manufacturer’s protocol. The staining reaction was stopped by adding FACS buffer (PBS containing 2% FBS, 2 mM EDTA, and 0.1% sodium azide) followed by centrifugation and washing. Cells were then incubated with conjugated antibodies at 1:100 dilution for 20 minutes at room temperature in the dark. Cells were subsequently washed thoroughly with ice-cold FACS buffer two additional times before acquisition. Data acquisition was performed using a FACSymphony A5 Cell Analyzer (BD Biosciences) equipped with 355, 405, 488, 532, and 640-nm lasers. Data analysis was conducted using FlowJo v10.10 software. Compensation beads for anti-mouse antibodies (BD, 552843) were used throughout the study for fluorescence compensation. For analysis of triple reporter activation in MART-1 TCR CD8⁺ Jurkat T cells co-cultured with artificial antigen-presenting cells (APCs), additional compensation beads for CFP, GFP, and mCherry (Thermo Fisher Scientific, A54742, A10514, A54743) were utilized.

### SLB-T cell interaction assays for ligand clustering and visualization

Supported lipid bilayers (SLBs) were generated using a previously established method (*78*) with minor modifications. Liposomes of small unilamellar vesicles (SUVs) for SLBs were prepared using a standard extrusion (Avanti, method following manufacturer’s protocol with a lipid composition of 91.75% 1,2-dioleoyl-sn-glycero-3-phosphocholine (DOPC, 850375), 6.25% DGS-NTA(Ni^2+^) (790404), and 2% Biotinyl Cap PE (870273). Clean room cleaned, high performance glass coverslips (SCHOTT Nexterion, 1472315) were assembled with flow channel chambers (Ibidi, 80608). Prepared 0.2 mM SUV liposomes were added to each chamber and incubated for 60 minutes at room temperature. After washing unbound liposomes, 12×histidine-tagged ligand molecules (fluorescently labeled or unlabeled) were incubated on the SLBs to attach to nickel lipids for 30 minutes at room temperature, followed by two washes. All main figure experiments contained immunological synapse formation with SLBs containing anti-TCR VHH-His (∼200 molecules/μm²) and ICAM-1-His (∼200 molecules/μm²). Additional ligand incorporation (CD80, CD86, or PD-L1) on SLBs is specified in each figure illustration, with a density of 200 molecules/μm² unless otherwise specified. Casein blocking buffer (*78*) was incubated for 20 minutes at 37°C and washed twice with HBS-BSA buffer (20 mM HEPES pH 7.2, 137 mM NaCl, 5 mM KCl, 0.7 mM Na₂HPO₄, 6 mM D-glucose, 2 mM MgCl₂, 1 mM CaCl₂, 1% w/v BSA) to minimize non-specific binding.

T cells were introduced into ligand-coated lipid bilayer chambers with or without blocking antibodies (5 μg/ml) where applicable. The final cell concentration was 1.0 × 10⁷ cells/ml for primary human CD3⁺ T cells, which were incubated for 40-45 minutes at 37°C, and 0.75 × 10⁷ cells/ml for Jurkat T cell lines (parental or CTLA4+), which were incubated for 30 minutes at 37°C otherwise depicted. Prior to spinning-disk confocal microscopy imaging, the SLB-T cell interface was fixed with 2% paraformaldehyde and washed thoroughly.

For soluble CTLA4-Ig assay, Jurkat T cells (0.75 × 10⁷ cells/ml) were introduced onto SLBs with recombinant CTLA4-Ig protein for selective visualization of CD28-B7 ligands interactions. T cells were incubated for 30 minutes at 37°C and fixed with 2% paraformaldehyde prior to spinning-disk confocal microscopy imaging.

For live cell imaging, stimulated human primary T cells (1.0 × 10⁷ cells/ml) were introduced onto SLBs. SLBs were pre-heated in the microscope chamber for at least 20 minutes prior to T cell injection. Immunological synapse formation was timed from the moment of T cell injection. At the 15-minute time point, either HBS-BSA buffer (control) or anti-PD-1 blocking antibody (5 μg/ml) was injected, and time-lapse imaging was initiated with 3-minute intervals.

### SLB - T cell interaction assays for visualization of synaptic T cell components using TIRFM

For PD-1 detection experiments, T cells were incubated on SLBs containing the appropriate ligand combinations as specified in figure legends. Following the standard incubation and fixation protocol, cells were then incubated with the non-competing, non-blocking anti-PD-1 scFv-AlexaFluor 647 at 100 nM at room temperature. TIRFM imaging was utilized to selectively illuminate and detect PD-1 molecules specifically at the cell-SLB interface, thereby eliminating background fluorescence from PD-1 molecules in the bulk cytoplasm and cell surface regions not in contact with the SLB.

For T cell adhesion assay with PKCθ staining to evaluate CD28 phosphorylation, fixed T cells on SLBs were permeabilized with 0.02% Triton-X-100 briefly and stained with 1:500 diluted anti-PKCθ primary antibody (Cell Signaling Technology, 86952) conjugated with AlexaFluor 488. Interference reflection microscopy (IRM) was performed to visualize T cell adhesion and contact areas with the SLB. Concurrent TIRFM imaging enabled specific detection of PKCθ recruitment to the immunological synapse, allowing for correlation between cell adhesion patterns and phosphorylated CD28 clustering.

### Microscopy set up

Imaging was performed on a Nikon CSU-W1 SoRa spinning disk confocal microscope equipped with a Hamamatsu sCMOS camera (Orca-Fusion-BT) and TIRF module. The microscopy system was equipped with 405, 488, 561, and 640-nm lasers. The spinning disk speed was set to a maximum of 4,000 rpm, which enabled visualization of microclusters of ligand molecules on supported lipid bilayers (SLBs). Imaging was conducted at room temperature for most experiments, except for live cell imaging, which was performed at 37°C to maintain physiological conditions. For spinning-disk confocal imaging using SoRa (super resolution by optical reassignment) mode, either a water-immersion 60× objective with 1× magnification or a 40× objective with 2.8× magnification was used. T cell adhesion was evaluated when needed using interference reflection microscopy (IRM) in epi-fluorescence mode. For total internal reflection fluorescence microscopy (TIRFM), an oil immersion 60x TIRF objective was used to achieve selective illumination of the cell-bilayer interface.

### Fluorescence recovery after photobleaching (FRAP) for fluidity test of SLBs

SLB fluidity and membrane integrity were assessed using fluorescence recovery after photobleaching (FRAP) assays. FRAP experiments were performed using the Leica SP5 confocal microscope with a 640-nm laser. A circular region of interest (ROI) with a diameter of 11 um was photobleached using 100% laser power for 5 frames. Fluorescence recovery was monitored by acquiring images every 1.29 seconds for 100.82 seconds post-bleaching. The mobile fraction (Plateau) was evaluated by fitting the exponential curve to the normalized FRAP data (Y).

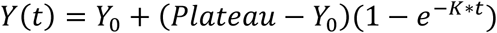

The normalized FRAP data (Y) was calculated by:

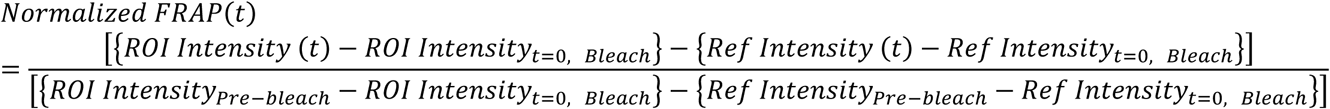

For more accurate diffusion coefficient calculation, *τ* was fitted to diffusion expression below (*79, 80*).

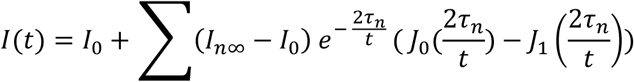

Then the diffusion coefficient was calculated with the expression below where *d* is a diameter at bleaching.

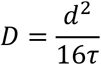

### Ligand density evaluation on SLBs using single-molecule imaging

Ligand density on SLBs was determined using total internal reflection fluorescence microscopy (TIRFM) on a Nikon CSU-W1 spinning disk confocal system equipped with an sCMOS camera. Fluorescence intensity measurements were performed to establish a quantitative relationship between fluorescence signal and molecular count, using a similar approach to previous literature (*81*). Briefly, the average fluorescence intensity of ligand molecules attached to SLBs was first recorded across the entire field of view (FOV), to establish the initial intensity. Controlled photobleaching was then applied to reduce fluorescent particle density to 0.2-0.3 per μm^2^ within FOV, ensuring spatial separation of individual ligand molecules. Further imaging captured resolvable proteins where each fluorescent spot represented a single ligand, irrespective of fluorophore conjugation stoichiometry. Automated single-molecule detection analysis provided the number of resolved ligand molecules, which was combined with average intensity across the FOV to generate a calibrated ‘intensity-to-count’ conversion factor. This calibration factor was subsequently applied to pre-bleach initial intensity to determine total ligand counts, with surface density calculated by normalizing to FOV area.

### Image analysis of immunological synapse

Analysis and evaluation of fluorescent signal from each immunological synapse were done by collecting intensity information of ligand molecules on SLBs or by collecting intensity information of PD-1 binding scFv at the immunological synapse. Image processing began with flatness correction to eliminate uneven illumination across the field of view. To account for dye-to-protein ratio variability and batch-to-batch variations in fluorescent protein preparations, intensity fold change parameter was analyzed to compare ligand interactions to receptors (e.g. comparison of CD80 and CD86 interactions to CD28). Intensity fold change analysis was performed by dividing the average intensity of the ligand-of-interest within the cell adhesion area to the average intensity of the ligand attached on SLBs outside the cell adhesion area. Otherwise, the average intensity of the ligand-of-interest within the cell adhesion area was used for analysis.

Interference reflection microscopy (IRM) signals were analyzed to distinguish well-adhered cells from poorly attached cells. Cells displaying IRM signals above defined thresholds for size, morphology, and signal intensity were classified as well-adhered. The number of well-adhered cells per field of view was counted and used for statistical analysis to assess the relationship between cell adhesion and experimental treatments.

### BSLB - T cell co culture Activation protocol

Bead-supported lipid bilayers (BSLBs) were generated using a previously established method (*78*) with minor modifications. Silica glass beads (5 µm, Bangs Laboratories SS05003) were used for BSLB preparation. Liposome solutions containing DOPC, DGS-NTA(Ni^2+^), and Biotinyl Cap PE at 91.75%, 6.25%, and 2% were mixed and washed thoroughly using HBS-BSA buffer (20 mM HEPES pH 7.2, 137 mM NaCl, 5 mM KCl, 0.7 mM Na₂HPO₄, 6 mM D-glucose, 2 mM MgCl₂, 1 mM CaCl₂, 1% w/v BSA). Beads were washed three times by centrifugation at 600xg, 2 min to remove unbound liposomes. Afterward, biotinylated MART-1-HLA complex was coupled with streptavidin at a 1:2 molar ratio, and 12×histidine-tagged ligands (ICAM-1, CD80 or CD86, with or without PD-L1) were incubated with BSLB-coated beads for 30 minutes at room temperature on an orbital plate shaker. BSLBs were thoroughly washed in HBS-BSA buffer two times. Finally, MART-1 TCR⁺ CD8⁺ Jurkat T cells with triple reporters were co-cultured with BSLBs at a 1:3 ratio to reach 1.0 × 10^6^ cells/ml of T cells and 3.0 × 10^6^ cells/ml of BSLBs for 18 hours in a 37°C, 5% CO₂ incubator. T cell activation was assessed by flow cytometric analysis of triple reporter expression (CFP, GFP, and mCherry) as indicators of NF-κB, NFAT, and AP-1. Cells were harvested and analyzed using the flow cytometry protocol described above.

### Statistical Analysis

Statistical analysis was performed using GraphPad Prism software (version 10.4.1). For comparisons between two groups, unpaired or paired Student’s t-tests were used, and for multiple group comparisons, one-way or two-way analysis of variance (ANOVA) was performed, followed by appropriate post-hoc tests as specified. Statistical significance was defined as P < 0.05, with specific P values indicated as follows: *P < 0.05, **P < 0.01, ***P < 0.001, ****P < 0.0001. The specific statistical tests used, number of independent experiments, and sample sizes are indicated in individual figure legends.

## Acknowledgments

We thank Karin Lee, Luca Melchiori and Discovery Immune Cell Engagers team in AstraZeneca Oncology R&D for support and thoughtful discussion on this study. We thank Scott Hammond and Mark Cobbold for the support of this project. We would like to acknowledge Jan Martinek of Microscopy Core Facility and Flow Cytometry Hub Team at Gaithersburg. Depictions are created with BioRender.com.

## Funding

This study was funded by AstraZeneca. G.E. is a fellow of the AstraZeneca Postdoctoral Research Program. M.L.D. is supported by the Kennedy Trust for Rheumatology Research and the Chinese Academy of Medical Sciences (CAMS) Innovation Fund for Medical Science (CIFMS), China (grant number: 2024-I2M-2-001-1).

## Author contributions

Conceptualization: GE, SJD, KNP, RV

Methodology: GE, VA, AG, MLD, RV

Investigation: GE, RV

Visualization: GE

Analysis: GE, JB

Supervision: KNP, MLD, RV

Writing-original draft: GE, RV

Writing-review and editing: GE, SC, SJD, KNP, MLD, RV

## Competing interests

GE, JB, VA, AG, SC, SJD, KNP and RV were employed by AstraZeneca at the time this work was performed.

## Data and materials availability

All data needed to evaluate the conclusions in the paper are present in the main text or the supplementary materials. The codes for the microscopy analysis in MATLAB will be deposited in GitHub.

## Notes

### Competing Interest Statement

GE, JB, VA, AG, SC, SJD, KNP and RV were employed by AstraZeneca at the time this work was performed.
This study was funded by AstraZeneca. G.E. is a fellow of the AstraZeneca Postdoctoral Research Program. M.L.D. is supported by the Kennedy Trust for Rheumatology Research and the Chinese Academy of Medical Sciences (CAMS) Innovation Fund for Medical Science (CIFMS), China (grant number: 2024-I2M-2-001-1).

