## Supplementary Figures for "PD-1 blockade enhances T cell activation by reorchestrating CD28 and CTLA4 ligand interactions"

Gee Sung Eun *et al.*

**This PDF file includes:**

Figs. S1 to S8
Table S1

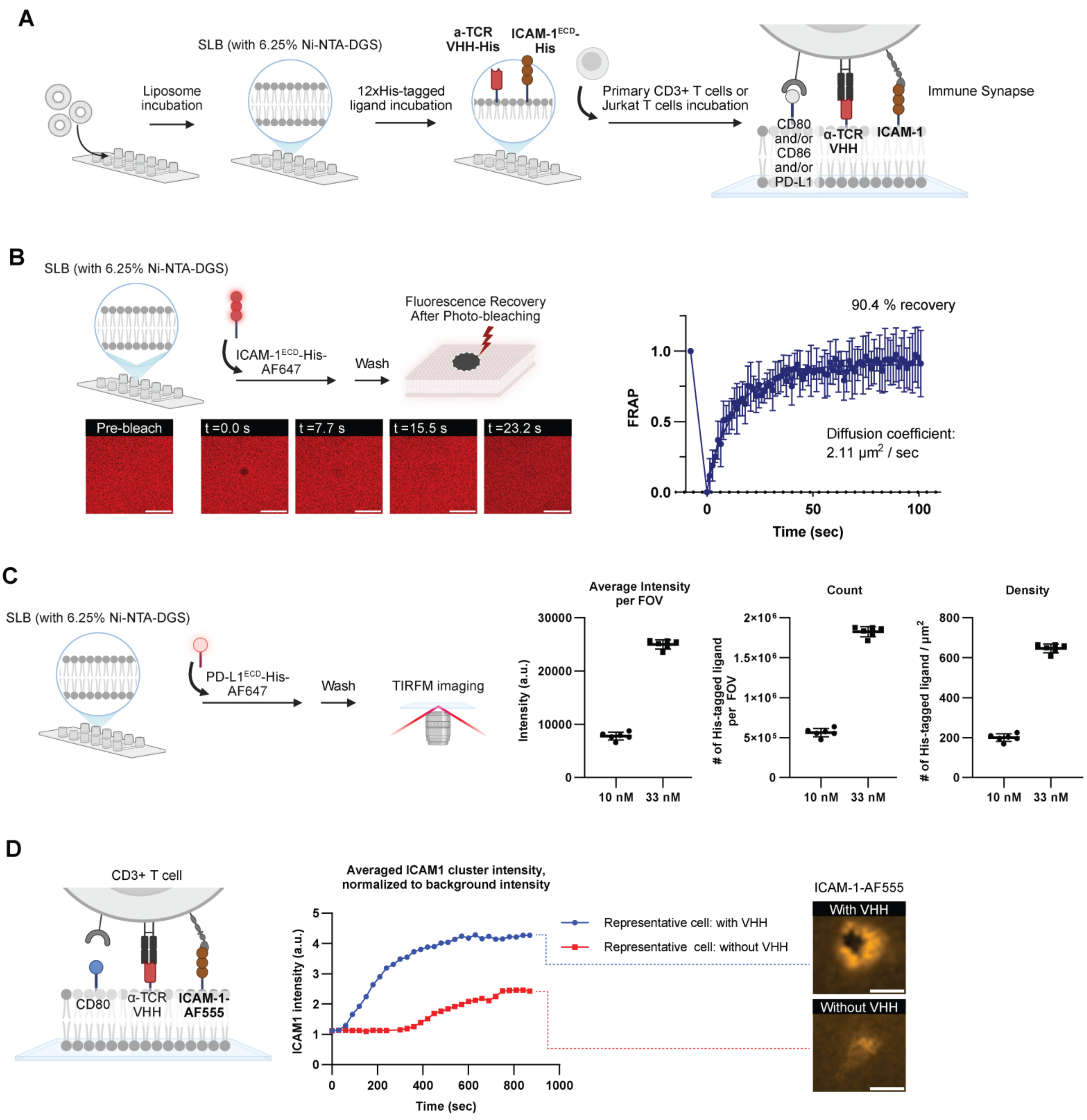


**Fig. S1.**
Characterization of supported lipid bilayers and immune synapse formation.

**(A)** Schematic of immune synapse formation. T cell and the artificial antigen-presenting cell (APC) interaction interface formed using supported lipid bilayers (SLBs) containing 6.25% $Ni^{2+}$-NTA-DGS lipids with anti-TCR VHH-His and ICAM1-His or various combinations of CD80, CD86 and PD-L1. All immune synapse experiments include anti-TCR VHH and ICAM-1 molecules on SLBs. **(B)** Fluorescence recovery after photobleaching (FRAP) experiment of SLBs containing AF647-labeled ICAM-1. Data are presented as mean ± S.D (N=8). Scale bars: 50 µm. **(C)** Characterization of histidine-tagged ligand molecule density on SLBs. Average intensity and molecule numbers counted per field of view (FOV) of ~2,826 $\mu m^2$. Data presented as mean ± S.D (N=6). Immune synapse experiments in our work typically contain ligand density of ~200 /$\mu m^2$ unless otherwise specified. **(D)** Freshly isolated human primary $CD3^+$ T cells forming immune synapse on bilayers containing ICAM-1, CD80, and -/+ anti-TCR VHH. Real-time live cell imaging was done to track AF555- labeled ICAM-1 clustering patterns. Control experiments without anti-TCR VHH molecule show no ring-like ICAM-1 patterns.

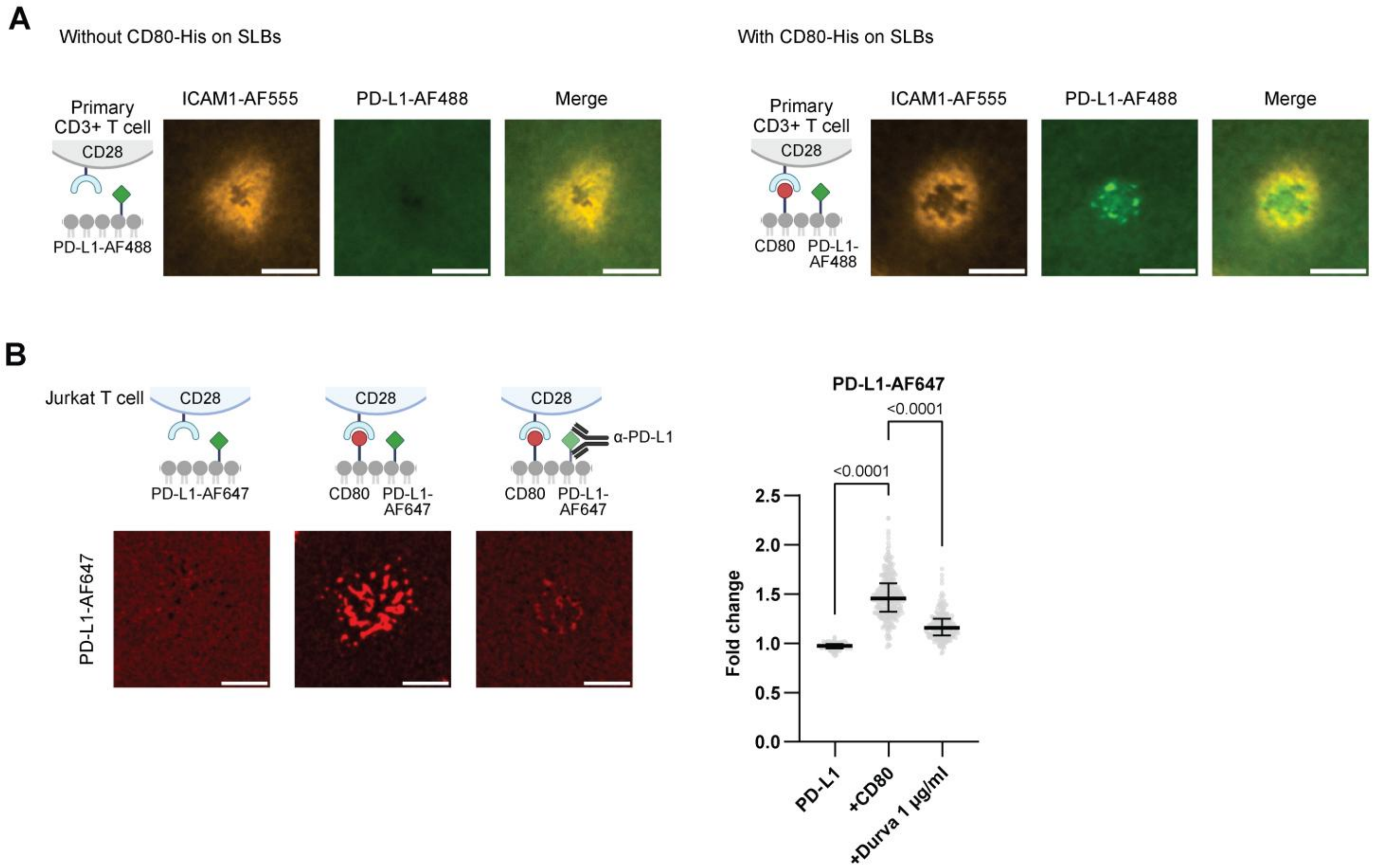


**Fig. S2.**

CD80-PD-L1 *cis*-interactions in immune synapses.

**(A)** Freshly isolated human primary $CD3^+$ T cells forming immune synapses on SLBs containing anti-TCR VHH, ICAM-1-AF555, PD-L1-AF488, with or without CD80. Left: PD-L1 clustering on SLB without CD80. Right: PD-L1 clustering on SLB with CD80. Scale bars: 5 µm. **(B)** Jurkat T cells forming immune synapses on SLBs containing anti-TCR VHH, ICAM-1, PD-L1-AF647, with or without CD80. PD-L1 blocking antibody (durvalumab) used in the third condition. Left: Schematic with representative images. Scale bars: 5 µm. Right: Quantification of PD-L1 accumulation per synapse as a fold change over area of bilayers with no cells. Data presented as column dot plots with median ± interquartile range ($N_{Cells} > 130$, from two independent experiments). Statistical analysis by one-way ANOVA.

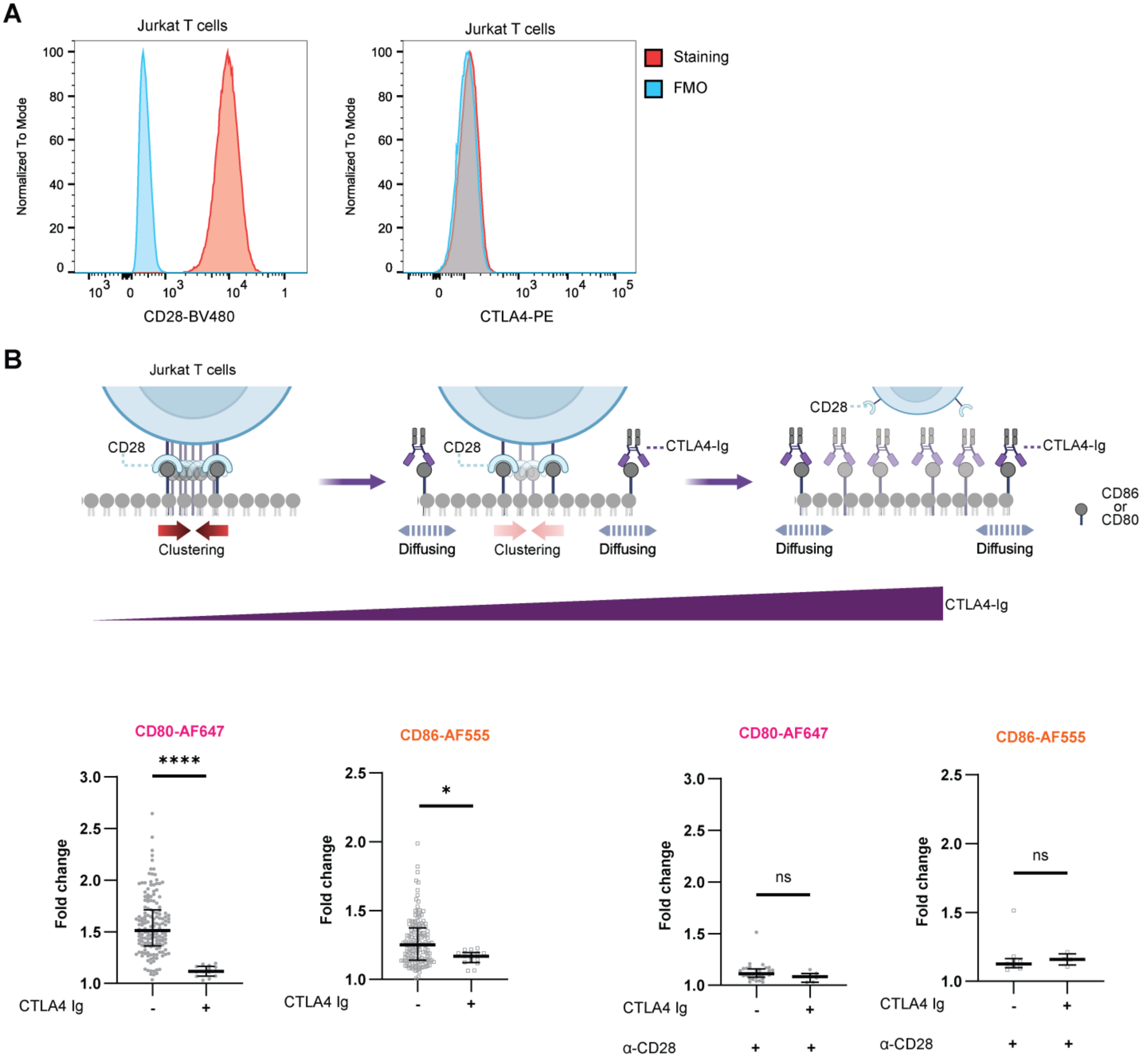


**Fig. S3.**

Validation of soluble CTLA4-Ig assay with Jurkat T cells that selectively visualizes CD28-ligand interactions.

**(A)** Jurkat T cells used in experiments here express CD28 but not CTLA4. **(B)** Upper panel: Schematic of Jurkat T cells forming immune synapses on SLBs containing anti-TCR VHH, ICAM-1, CD80 or CD86. Soluble CTLA4-Ig binds to CD80 or CD86 on SLB without inducing ligand clusters. Ligand clustering results exclusively from CD28 interactions. Lower panel: Quantification of CD80-AF647 or CD86-AF555 accumulation per synapse as a fold change over area of bilayer with or without CTLA4-Ig, including anti-CD28 blockade controls. Data presented as column dot plots with median ± interquartile range. *, $p<0.05$; ****, $p<0.0001$; Statistical analysis by unpaired t-test.

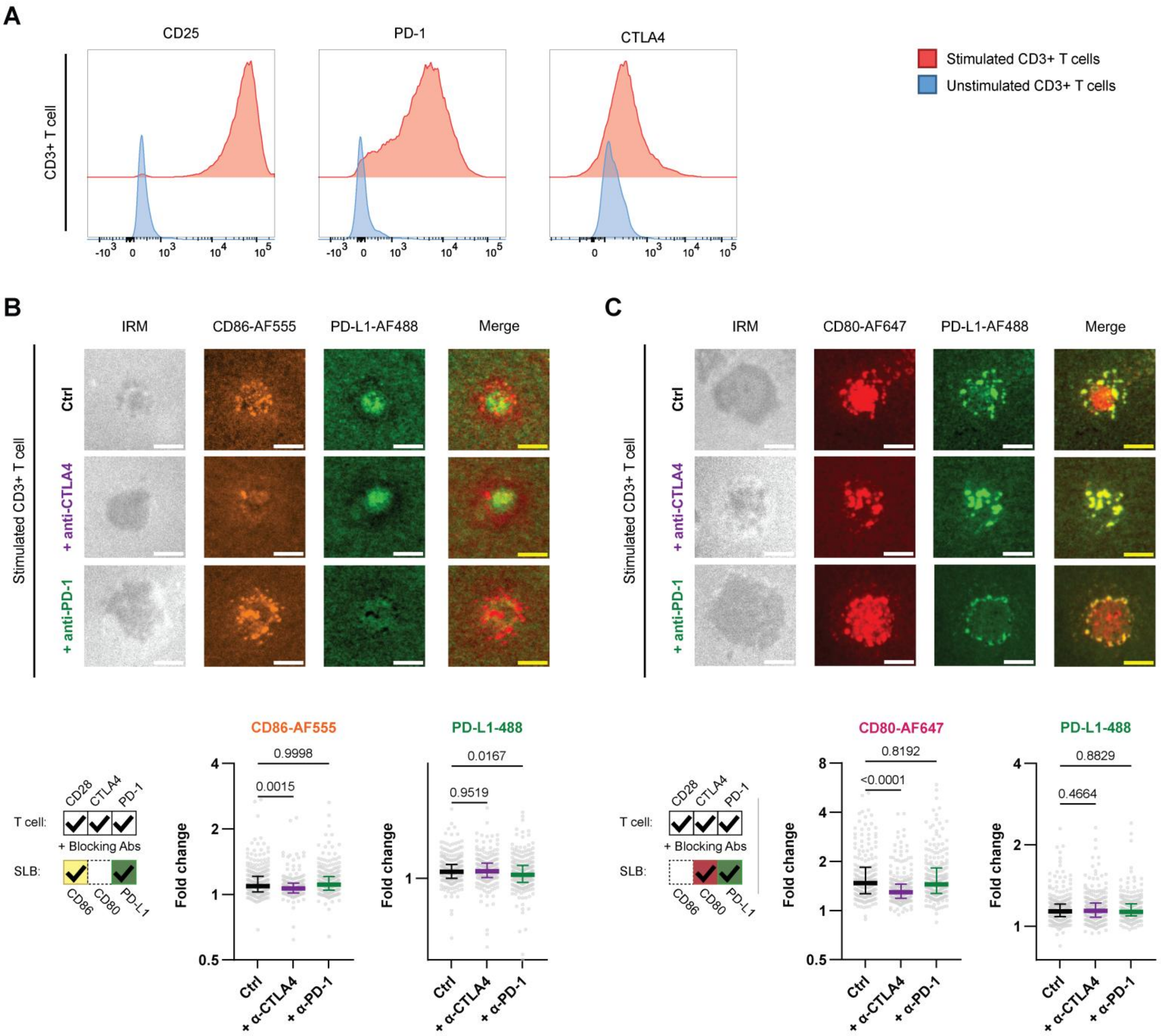


**Fig. S4.**

Checkpoint blockade effects on CD86/PD-L1 or CD80/PD-L1 ligand accumulation in stimulated T cells.

**(A)** Expression of CD25, PD-1, and CTLA4 in stimulated and unstimulated human primary $CD3^+$ T cells. **(B)** Stimulated human primary $CD3^+$ T cells forming immune synapses on SLBs containing anti-TCR VHH, ICAM-1, CD86-AF555, and PD-L1-AF488 with or without blocking antibodies. Upper: Representative IRM images and ligand accumulation of CD86-AF555, and

PD-L1-AF488. Scale bars: 5 µm. Lower: Quantification of CD86 and PD-L1 accumulation per synapse as a fold change over area of bilayer with no cell. **(C)** Stimulated human primary $CD3^+$ T cells forming immune synapses on SLBs containing anti-TCR VHH, ICAM-1, CD80-AF647, and PD-L1-AF488 with or without blocking antibodies. Upper: Representative IRM images and ligand accumulation of CD80-AF647, and PD-L1-AF488. Scale bars: 5 µm. Lower: Quantification of CD80, and PD-L1 accumulation per synapse as a fold change over area of bilayer with no cell. Data presented as column dot plots with median ± interquartile range ($N_{Cells}$ > 200 from three independent experiments). P values determined by one-way ANOVA with multiple comparisons using Dunnett's correction.

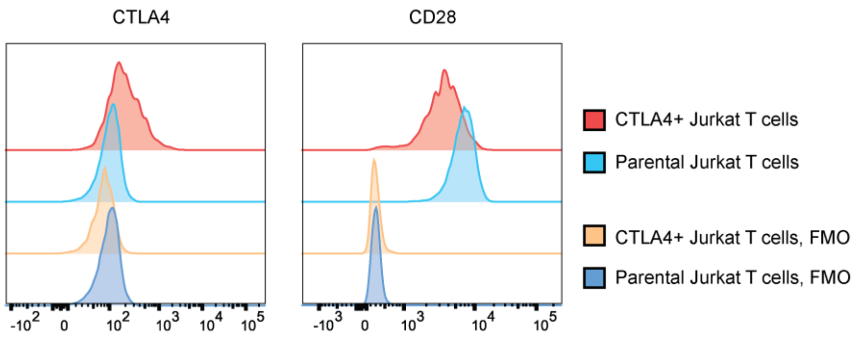


**Fig. S5.**

Expression of CTLA4 and CD28 in CTLA4+ Jurkat T cells. Fluorescence minus one (FMO) control shown in parallel.

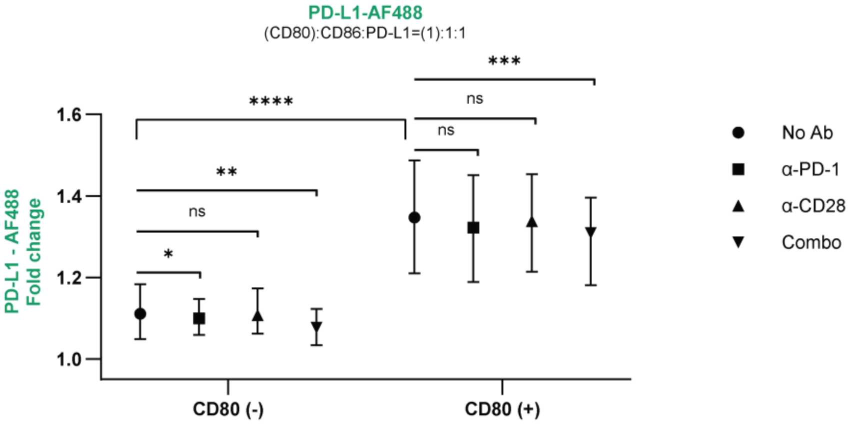


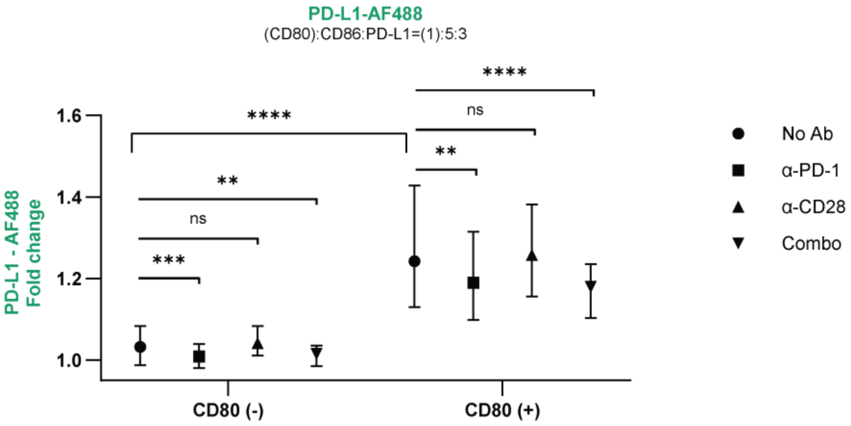


**Fig. S6.**

Quantification of PD-L1 accumulation in immune synapses from the experiment in Fig. 5. Stimulated primary $CD3^+$ T cells were incubated on SLBs containing anti-TCR VHH, ICAM-1 and various combinations of CD86, PD-L1-AF488, and -/+ CD80 as shown in the schematic and imaged using total internal reflection fluorescence microscopy (TIRFM) in the presence of non-blocking and non-competing AF647-labeled anti-PD-1 scFv to visualize PD-1. Quantification of PD-L1-AF488 accumulation per synapse as a fold over area of bilayer with no cell. Data is presented as column dot plots with median ± interquartile range ($N_{Cells}$ >120 from three independent experiments per condition). *, $p<0.05$; **, $p<0.01$; ***, $p<0.001$; ****, $p<0.0001$; P values determined by one-way ANOVA.

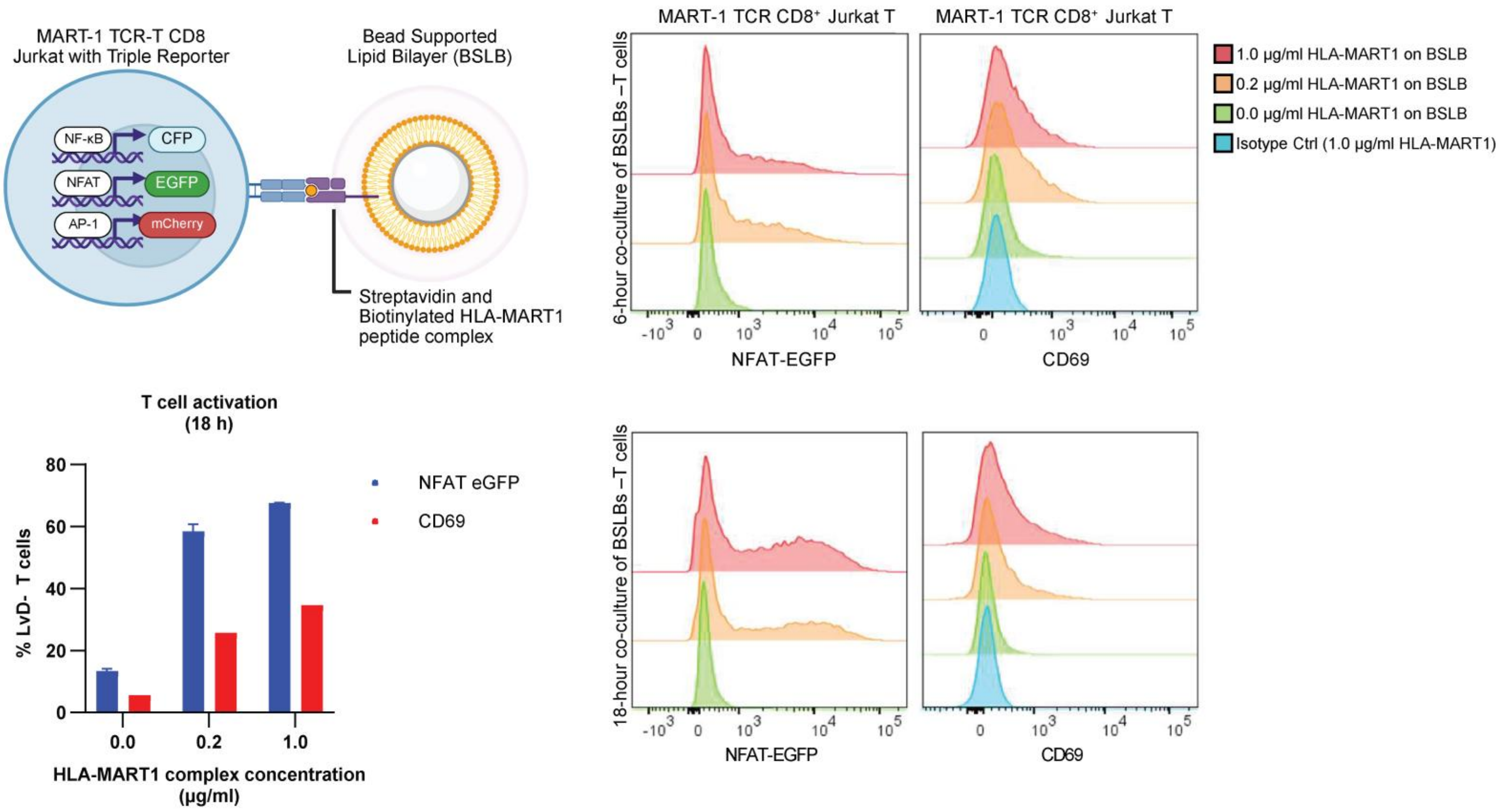


**Fig. S7.**

Characterization of MART-1 antigen-specific CD8$^+$ triple reporter Jurkat T cells for BSLB co-culture assay.

T cell activation was demonstrated at two doses of MART1 antigen and evaluated using NFAT-eGFP and CD69 expression at two different time points.

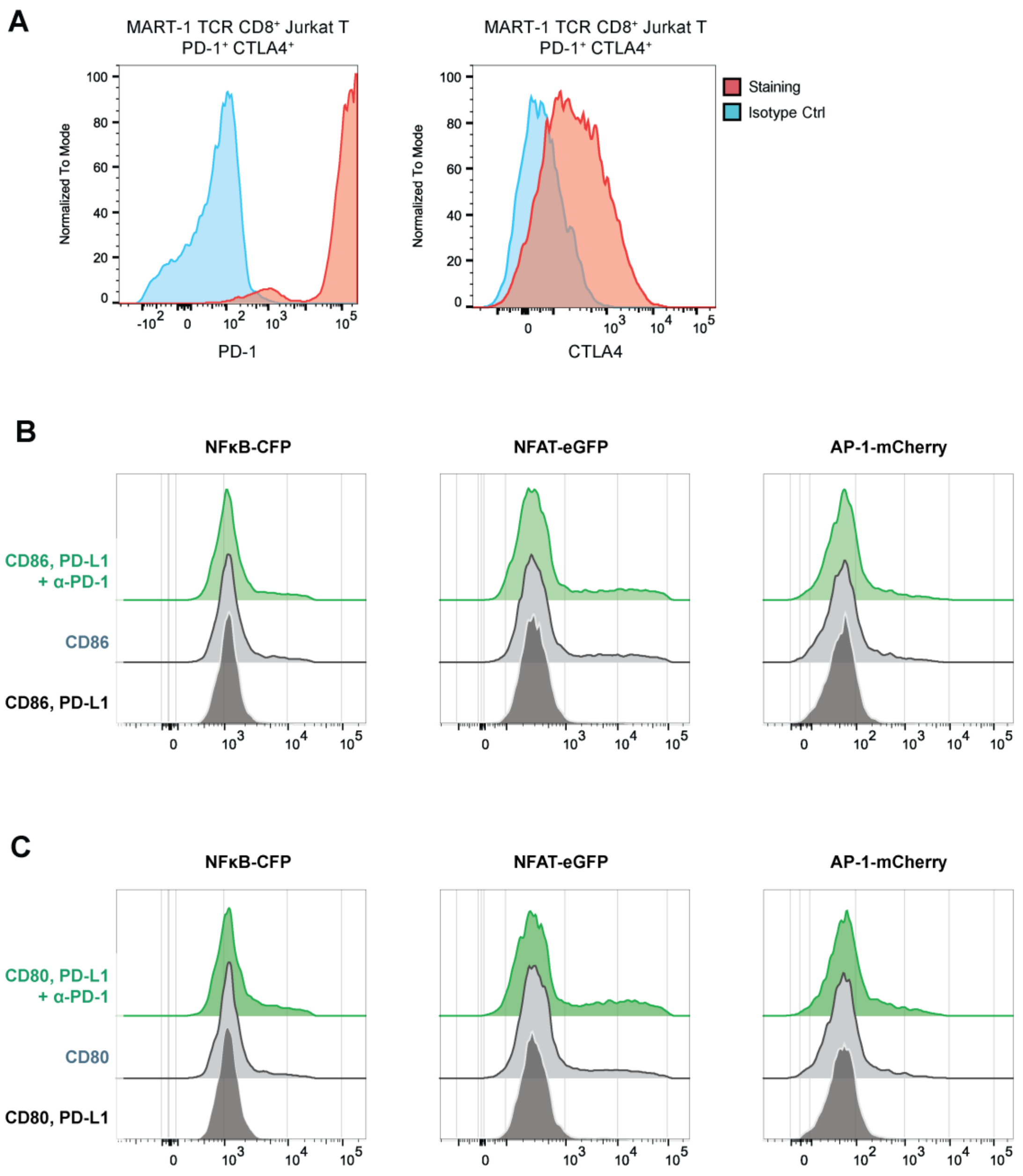


**Fig. S8.**

Co-culture experiments with CTLA4+ PD-1+ MART-1 TCR Jurkat cells and bead-supported lipid bilayers (BSLBs).

**(A)** PD-1 and CTLA4 expression in MART-1 TCR CD8$^{+}$ triple reporter Jurkat T cells transduced with PD-1 and CTLA4. **(B-C)** Histograms of T cell activation marker expression analyzed by flow cytometry under conditions containing CD86, CD86+PD-L1 and CD86+PD-L1+anti-PD-1 **(B)** or CD80, CD80+PD-L1 and CD80+PD-L1+anti-PD-1 **(C)** showing that PD-1 blockade enhances T cell activation by promoting CD28-CD80-PD-L1 interactions.

**Table S1.**

Recombinant Proteins

| Recombinant Protein | Sequence |
|---|---|
| ICAM-1-His | MGWSCIILFLVATATGVHSGNAQTSVSPSKVILPRGGSVLVTCSTSCDQPKLLGIETPLPKK<br>ELLLPGNNRKVYELSNVQEDSQPMCYSNCP<br>DGQSTAKTFLTVYWTPERVELAPLPSWQPVGKNLTLRCQVEGGAPRANLTVVLLRGEKE<br>LKREPAVGEPAEVTTTVLVRRDHHGANFSCR<br>TELDLRPQGLELFENTSAPYQLQTFVLPATPPQLVSPRVLEVDTQGTVVCSLDGLFPVSEA<br>QVHLALGDQRLNPTVTYGNDSFSAKASVSV<br>TAEDEGTQRLTCAVILGNQSQETLQTVTIYSFPAPNVILTKPEVSEGTEVTVKCEAHPRAK<br>VTLNGVPAQPLGPRAQLLLKATPEDNGRSFS<br>CSATLEVAGQLIHKNQTRELRVLYGPRLDERDCPGNWTWPENSQQTPMCQAWGNPLPEL<br>KCLKDGTFPLPIGESVTVTRDLEGTYLCRAR<br>STQGEVTRKVTVNVLSPRYEHHHHHHHHHHHH* |
| CD80-His | MGWSCIILFLVATATGVHSVIHVTKEVKEVATLSCGHNVSVEELAQTRIYWQKEKKMVL<br>TMMSGDMNIWPEYKNRTIFDITNNLSIVILALRPS<br>DEGTYECVVLKYEKDAFKREHLAEVTLSVKADFPTPSISDFEIPTSNIRRIICSTSGGFPEPH<br>LSWLENGEELNAINTTVSQDPETELYAVSSKL<br>DFNMTTNHSFMCLIKYGHLRVNQTFNWNTTKQEHFPDNLLPSHHHHHHHHHHHH* |
| CD80-KKCK-His | MGWSCIILFLVATATGVHSVIHVTKEVKEVATLSCGHNVSVEELAQTRIYWQKEKKMVL<br>TMMSGDMNIWPEYKNRTIFDITNNLSIVILALRPS<br>DEGTYECVVLKYEKDAFKREHLAEVTLSVKADFPTPSISDFEIPTSNIRRIICSTSGGFPEPH<br>LSWLENGEELNAINTTVSQDPETELYAVSSKL<br>DFNMTTNHSFMCLIKYGHLRVNQTFNWNTTKQEHFPDNLLPSKKCKGGSHHHHHHHHH<br>HHH* |
| CD86-His | MGWSCIILFLVATATGVHSLKIQAYFNETADLPCQFANSQNQSLSELVVFWQDQENLVLN<br>EVYLGKEKFDSVHSKYMGRTSFDSDSWTLRL<br>HNLQIKDKGLYQCIIHHKKPTGMIRIHQMNSELSVLANFSQPEIVPISNITENVYINLTCSSIH<br>GYPEPKKMSVLLRTKNSTIEYDGVMQKSQDN<br>VTELYDVSISLSVSFPDVTSNMTIFCILETDKTRLLSSPFSIELEDPQPPPDHIPHHHHHHHH<br>HHHH* |

| | |
|---|---|
| PD-L1-His | MGWSCIILFLVATATGVHSFTVTVPKDLYVVEYGSNMTIECKFPVEKQLDLAALIVYWEM EDKNIIQFVHGEEDLKVQHSSYRQRARLLKDQL SLGNAALQITDVKLQDAGVYRCMISYGGADYKRITVKVNAPYNKINQRILVVDPVTSEHE LTCQAEGYPKAEVIWTSSDHQVLSGKTTTTNSK REEKLFNVTSTLRINTTTNEIFYCTFRRLDPEENHTAELVIPELPLAHPPNERHHHHHHHH HHH* |
| Anti-TCR VHH-His | MGWSCIILFLVATATGVHSEVQLVESGGGLVQPGGSLRLSCVASGDVHKINFLGWYRQA PGKEREKVAHISIGDQTDYADSAKGRFTISRDESKNMVYLQMNSLKPEDTAVYFCRAFSR IYPYDYWGQGTLVTVSSHHHHHHHHHHHH* |
| PD-1 scFv | MGWSCIILFLVATATGVHSSYELTQDPAVSVALGQTVRITCSGGSSDYYGWFQQKPGQAP VTVIYYNNKRPSDIPDRFSGSSSGNTASLTIT GAQAEDEADYYCGNADSSVGVFGSGTKVTVLGKPGSGKPGSGKPGSGKPGSEVQLLESG GGLVQPGGSLRLSCAASGFTFSSYNMFWV RQAPGKGLEFVAEISGSNTGSRTWYAPAVKGRATISRDNSKNTLYLQMNSLRAEDTAVY YCAKSIYGGYCAGGYSCGVGLIDAWGQGTLV TVSSDYKDDDDKGGGSRGVPHIVMVDAYKRYK* |
